# Functional implications of the interaction of the SARS-CoV-2 Nucleocapsid protein with factors involved in nonsense-mediated mRNA decay

**DOI:** 10.1101/2024.07.16.603698

**Authors:** Megha Mallick, Volker Boehm, Guangpu Xue, Mark Blackstone, Niels H. Gehring, Sutapa Chakrabarti

## Abstract

The RNA genome of the SARS-CoV-2 virus encodes for four structural proteins, 16 non-structural proteins and nine putative accessory factors. A high throughput analysis of interactions between human and SARS-CoV-2 proteins identified multiple interactions of the structural Nucleocapsid (N) protein with RNA processing factors. The N-protein, which is responsible for packaging of the viral genomic RNA was found to interact with two RNA helicases, UPF1 and MOV10 that are involved in nonsense-mediated mRNA decay (NMD). Using a combination of biochemical and biophysical methods, we investigated the interaction of the SARS-CoV-2 N-protein with NMD factors at a molecular level. Our studies led us to identify the core NMD factor, UPF2, as an interactor of N. The viral N-protein engages UPF2 in multipartite interactions and can negate the stimulatory effect of UPF2 on UPF1 catalytic activity. N also inhibits UPF1 ATPase and unwinding activities by competing in binding to the RNA substrate. We further investigate the functional implications of inhibition of UPF1 catalytic activity by N in mammalian cells. The interplay of SARS-CoV-2 N with human UPF1 and UPF2 does not affect decay of host cell NMD targets but might play a role in stabilizing the viral RNA genome.

## Introduction

Coronaviruses belong to the order *Nidovirales*, which derives its name from the “nested”- subgenomic (sg) mRNA (similar 5’-ends and identical 3’-ends) transcribed upon infection of host cells [1, 2]. Approximately two-thirds of the 5’-end of the RNA genome of severe acute respiratory syndrome-related (SARS) coronaviruses comprise open reading frames (ORFs) 1a and 1b that are related by a 1 nucleotide ribosomal frameshift. They encode for two large polyproteins that are proteolytically processed to yield 16 non-structural proteins (Nsps) [3, 4]. These Nsps together form the replication-transcription complex (RTC) which is necessary for viral RNA synthesis [5, 6]. Discontinuous transcription of the 3’-end of the RNA genome leads to production of negative-sense sgRNAs that serve as templates for nested positive-strand sg-mRNAs, which can be translated to produce 4 structural proteins (Spike, Envelope, Membrane and Nucleocapsid, also referred to as S, E, M and N, respectively) and multiple accessory proteins [7, 8]. Continuous transcription from the 3’-end leads to synthesis of a full- length negative-sense RNA, which is used as a template for generation of new positive-sense genomic RNA. The viral genomic RNA is packaged by the Nucleocapsid (N) protein and enveloped by the remaining structural proteins to produce a helical nucleocapsid, in contrast to icosahedral nucleocapsids common to other positive-strand RNA viruses [9, 10].

The sg-mRNAs resemble host cell mRNAs in that they are capped at the 5’-end and polyadenylated at the 3’-end, enabling recognition of these mRNAs by the host cell translation machinery [11]. Although the 5’-leader sequence is identical in all sg-mRNAs, the lengths of the 3’-untranslated regions (3’-UTRs) greatly differ, ranging from 100-4000 nucleotides, depending on the site of initiation of discontinuous transcription. The genomic RNA, the 5’-end of which serves as a template for translation of the two large polyproteins, has the longest 3’-UTR of ∼10 kb [12]. In eukaryotes, transcripts with long 3’-UTRs are known to potentially trigger the nonsense-mediated mRNA decay (NMD) pathway [13–15]. The NMD pathway is a translation-dependent mechanism to detect and degrade mRNAs with unusual features such as premature termination codons (PTCs) or long 3’-UTRs [16, 17]. Certain sg-mRNAs as well as the genomic RNA of coronaviruses are, therefore, putative NMD targets. Interestingly, a study by Wada and co-workers using the murine hepatitis virus (MHV) as a model coronavirus reported an intricate interplay of the MHV-N protein with the NMD pathway at different stages of infection [18]. At an early stage of infection, when levels of N are low, viral replication was shown to be impeded by NMD. Upon sufficient accumulation of N in the later stages of infection, NMD is successfully inhibited by the N-protein, allowing for efficient viral replication. A recent study by Gordon and co-workers on mapping the SARS-CoV-2 – human interactome revealed interactions between the SARS-CoV-2 N-protein (hereafter referred to as N) and the RNA helicases, UPF1 and MOV10, which are known to play important roles in the NMD pathway [19].

The RNA helicase UPF1 (***Up***-***f***rameshift protein *1*) is a central component of the NMD pathway and along with the core NMD factors UPF2 and UPF3, is conserved from yeast to humans [20–22]. While the role of UPF1 in NMD was primarily thought to be to remodel nonsense mRNPs prior to degradation, it now appears that the RNA-dependent ATPase activity of UPF1 is also essential for discerning between cognate and non-cognate NMD substrates [23, 24]. Furthermore, UPF1 also serves as a phosphorylation-dependent protein-protein interaction platform to recruit factors that are important for progression of NMD and ultimately, degradation of the target mRNA [25–28]. The low basal catalytic activity of UPF1 is greatly stimulated upon binding to UPF2, which induces a large conformational change that switches the helicase from an inactive RNA-clamping mode to an active RNA-unwinding mode [29, 30]. Recently, it was shown that UPF2 triggers release of UPF1 from RNA, suggesting a rapid activation-and-dissociation mode of action of the helicase and its activator [31]. Although UPF1 is a key player of the NMD pathway, it is not the only RNA helicase involved. The helicases DHX34 and MOV10 were also shown to be involved in the NMD pathway, albeit at very different stages. DHX34 is thought function as a scaffold to recruit UPF1 to the PI3K-like kinase SMG1 to mediate its phosphorylation while MOV10 acts at a later stage of NMD, presumably to unwind RNA secondary structures in the 3’-UTR and facilitate decay [32, 33]. The observed interaction of N with UPF1 and MOV10 as determined by affinity-purification mass-spectrometry (AP-MS) suggest a direct perturbation of the NMD pathway by the N-protein and corroborates previous observations on the impact of MHV-N on NMD [19]. To investigate the molecular mechanisms of the interplay of coronavirus N-proteins and NMD, we adopted a biochemical approach to systematically test the interaction of SARS-CoV-2 N with the NMD factors, UPF1, UPF2 and MOV10. Surprisingly, we found that the N-protein does not bind UPF1 or MOV10 directly but instead interacts with UPF2. Using further biochemical and biophysical assays, we showed that N engages UPF2 in multi-partite interactions and indirectly associates with UPF1 to repress its catalytic activity. However, inhibition of UPF1 activity does not translate to inhibition of host cell NMD by N. We speculate that the N-protein might specifically inhibit NMD of viral genomic and sub-genomic mRNA while leaving host-cell NMD unaffected.

## Materials and Methods

### Protein expression and purification

All human UPF1 (except for full-length UPF1) and UPF2 constructs as well as SARS-CoV-2 N variants used in this study were expressed in *Escherichia coli* as 6X-His or His-GST fusions that are cleavable by TEV or HRV-3C protease. Details of the different constructs used in this study are provided in Table S1. The primers used to generate the different N expression constructs are described in Table S2. The SARS-CoV-2 Nfl plasmid was a gift from Markus Wahl. All plasmids produced for this study were verified by Sanger sequencing. Protein expression in *Escherichia coli* BL21(DE3) star pRare cells was induced by addition of isopropyl β-D-1-thiogalactopyranoside (IPTG) and performed at 18 °C for 20 hours. Full-length UPF1 was expressed using a baculovirus expression system (see Supplementary methods). All proteins were first subjected to Nickel-affinity chromatography, followed by Heparin-affinity chromatography and size-exclusion. Briefly, cells expressing the desired protein were resuspended in lysis buffer A (50 mM Tris-HCl pH 7.5, 500 mM NaCl, 10 mM imidazole and 10 % glycerol) supplemented with 1 mM phenylmethylsulphonyl fluoride (PMSF) and 0.5 mg of DNase I. All buffers used for purification of UPF1 and MOV10fus were further supplemented with 1 mM MgCl2, 1 µM ZnCl2 and 0.1 M urea. The cell suspension was lysed by sonication, following which the lysate containing soluble protein was separated from the cell debris by centrifugation and filtration. The clarified cell lysate was incubated with Ni^2+^-affinity resin (Machery-Nagel #745400.100). His-tagged protein was eluted in elution buffer B (20 mM Tris-HCl pH 7.5, 150 mM NaCl, 300 mM imidazole and 10 % glycerol). The one-step purified protein was then loaded onto a HiTrap Heparin Sepharose HP column (GE Healthcare) and eluted from it in a salt gradient of 100-1000 mM NaCl. Size exclusion chromatography (SEC) using Superdex 75 or Superdex 200 columns (GE Healthcare) was performed in SEC buffer C (20 mM Tris-HCl pH 7.5, 150 mM NaCl, 5 % glycerol and 2 mM DTT) as a final step to ensure homogeneity of the isolated protein. For all N-proteins, the pH of the SEC buffer was maintained at 8.0. In every case, the purity of the target protein was verified by SDS-PAGE after each chromatographic step.

To purify the UPF1-UPF2 and UPF2-UPF3 complexes, proteins were mixed in a 1 to 1.2 molar ratio (with UPF1 in excess for the UPF1-UPF2 complex, and UPF3 in excess for the UPF2- UPF3 complex) for 16 hours at 4 °C and resolved on a Superdex 200 column using SEC buffer C.

### GST pull-down assays

8 µg of bait and prey proteins were mixed and diluted to 40 µl in GST pulldown buffer (20 mM HEPES pH 7.5, 150 mM NaCl, 10 % glycerol and 0.1 % NP-40). 0.4 µg of the protein mixture was used as an input for each reaction. The protein mixtures were incubated on ice for 1 hour. following which 12 µl of a 50 % Glutathione Sepharose resin (GE Healthcare) was added to each sample. The mixture was further supplemented with 150 µl of GST-pulldown buffer and incubated at 4 °C for 90 minutes. The beads were extensively washed with same buffer before eluting the bound proteins in GST-pulldown buffer supplemented with 20 mM glutathione. The inputs and elutions were analysed by SDS-PAGE and Coomassie staining.

### Isothermal titration calorimetry (ITC)

ITC analyses were conducted to determine the binding affinities of UPF2 constructs (UPF2S, UPF2-MIF4G3, and UPF2-U1BD) and the NIDR1-core protein at 10 °C on an iTC200 instrument (MicroCal). All proteins were re-purified by SEC (Superdex 200 10/300) using a SEC/ITC buffer (20mM Tris-HCl, pH 8.0, 150 mM NaCl and 5% glycerol) immediately before their use in ITC. 40 µl of 200 µM of each UPF2 protein was loaded in the syringe and titrated against 250 µl of 20 µM NIDR1-core in a series of injections. 20 injections were performed in total, where the first injection was of 0.5 µl, followed by 19 injections of 2 µl each at 180-second intervals. Analysis of Ncore binding to UPF2S was carried out using an identical procedure. The data were analysed with MicroCal PEAQ-ITC Analysis Software (Malvern Instruments) by subtracting baseline and offset. All N variants were considered a dimer for concentration measurements and data analyses in all experiments requiring molar concentrations of proteins.

### Analytical size exclusion (SEC)

1000 pmol of the UPF1-UPF2 and UPF2-UPF3 complexes were mixed with 1000 pmol of N-proteins to a final volume of 50 µl in a SEC buffer D (20 mM HEPES pH 7.5, 150 mM NaCl, 2 % glycerol, 1 mM MgCl2, 1 µM ZnCl2, 2 mM DTT. Similarly, for a 1:1 and 1:2 UPF1-RNA-N complex, 500 pmol of UPF1, 500 pmol of U45 RNA and 500 pmol (for 1:1 mixture) or 1000 pmol (for 1:2 mixture) of NΔL were mixed and incubated at room temperature for 1 hour. The protein or protein-RNA mixtures were resolved on a Superdex 200 Increase 5/150 GL column in SEC buffer D. The peak fractions were analysed by SDS-PAGE and Coomassie staining to visualize proteins. To detect RNA, 3 µl of each peak fraction was 5’-end labelled with [γ-^32^P]- ATP using T4 polynucleotide kinase. The labelling reactions were resolved on a 12 % Urea-PAGE gel and visualized by phosphorimaging.

### Fluorescence-based unwinding assay

The sequences of the RNA and DNA strands of the RNA:DNA hybrid and that of the trap DNA strand are as follows:

RNA: 5’-GGGACACAAAACAAAAGACAAAAACACAAAACAAAAGACAAAAACACAAAACAAAAGACAAAAAGCCAAAUUACCGUGUGCGUACAACUAGCU-3’

Complementary DNA strand: 5’-GTGTGCGTACAACTAGCT-3’

Trap: 5’-GTGTGCGTACAACTAGC-3’

The DNA strand of the RNA:DNA hybrid was labelled at the 5’-end with Alexa Fluor 488 and the complementary trap DNA strand was labelled at the 3’-end with a quencher (Blackhole quencher 1).

The RNA substrate used in this helicase assay was prepared by *in vitro* transcription (IVT) from a linearised double-stranded (ds) DNA template. The DNA oligos used as the template for IVT are described in Table S2. The RNA:DNA duplex substrate for the unwinding assay was freshly prepared prior to every set of experiments by incubating the DNA and RNA strands in a 7:11 ratio with 2 mM MgOAc and 1x unwinding buffer (10 mM MES pH 6.5, 50 mM KOAc, 0.1 mM EDTA) at 95 °C for 210 seconds, followed by slow cooling of the mixture down to 30 °C. For each replicate, an initial reaction mixture was prepared which included 75 nM of freshly assembled RNA-DNA duplex in 1x unwinding buffer, 2 mM magnesium acetate, 2 mM freshly prepared DTT, and 300 nM UPF1, 600 nM UPF2 wherever mentioned, and 300- 1200 nM of Nfl. The reaction mixture containing UPF1 or UPF1-UPF2 was incubated at 25 °C for 10 minutes prior to adding N. The mixture was incubated for another 10 minutes after addition of N. Trap DNA was added at the very end to a final concentration of 0.56 µM. The reaction mixture was prepared in the dark and transferred to a 384-well plate (PerkinElmer OptiPlate 384-F). 2 mM ATP was injected into each well using the injector module of the Spark multimode microplate reader (Tecan Life Sciences). The fluorescence was monitored for 30 minutes at 30 °C on the same instrument. The measured fluorescence intensities were normalized to the 0-time point (baseline) for each condition to obtain relative fluorescence. A more detailed protocol in the form of a flow chart can be found in Supplementary methods.

### ATPase assay

The ATPase activity of UPF1 alone and in presence of its activator UPF2, and different amounts of Nfl was determined by quantifying the amount of inorganic phosphate released upon ATP hydrolysis using a coupled colorimetric assay (EnzCheck Phosphate Kit, Thermo Fischer Scientific). The protein mixtures were pre-incubated in a reaction volume of 150 µL with 1 µg poly-U-RNA, 40 nmol MESG (2-amino-6-mercapto-7-methylpurine ribonucleoside), and 0.5 U purine-nucleoside phosphorylase in an ATPase reaction buffer (50 mM MES pH 5.5, 50 mM potassium acetate, 5 mM magnesium acetate and 2 mM DTT) at 30 °C for 20 minutes. The reaction was initiated by addition of 1 mM ATP (final concentration in a final volume of 200 µL) in a 96-well plate (Greiner). 2-amino-6-mercapto-7-methylpurine produced upon cleavage of MESG by the inorganic phosphate generated from the ATPase reaction was detected by measuring absorbance at 360 nm on a Spark multimode microplate reader (Tecan Life Sciences). The reaction was monitored over a period of 20 minutes at 60-second intervals. The amount of UPF1 was kept constant (i.e., 6 pmol or final concentration of 200 nM) in each experiment, and UPF2 was added in 2-fold molar excess of UPF1 (12 pmol or final concentration of 400 nM). Nfl was added as indicated in the figures. The end point of the ATPase reaction (15-minute time point) of the UPF1-UPF2S mixture in each case was set to 1 or 100 %, and all other values were normalized to this maximum value.

### Cell culture

Flp-In T-REx-293 (human, female, embryonic kidney, epithelial; Thermo Fisher Scientific, RRID:CVCL_U427) cells were cultured in high-glucose, GlutaMAX DMEM (Gibco) supplemented with 9% fetal bovine serum (Gibco) and 1x Penicillin Streptomycin (Gibco). The cells were cultivated at 37°C and 5% CO2 in a humidified incubator. Stable cell lines were generated by using the PiggyBac (PB) Transposon system with the cumate-inducible PB-CuO-MCS-BGH-EF1-CymR-Puro vector. This vector was modified from the original vector (PB-CuO-MCS-IRES-GFP-EF1α-CymR-Puro (System Biosciences)) by replacing the IRES-GFP cassette with a BGH polyA signal. Inserts coding for N proteins were cloned either from IDT 2019-nCoV_N_Positive Control (Catalog # 10006625, CoV-2-N (bat)) or from SARS-CoV- 2 Nfl plasmid (mentioned above, CoV-2-N (human)). Transfection of cells was performed using a calcium phosphate-based system with BES buffered saline (BBS). Stably integrated cells were selected using media containing 2 μg/ml puromycin (InvivoGen) for a week. To express the N-terminally FLAG-tagged protein constructs, 2.8 x 10^5^ cells were seeded in 6-well plates, 30 µg/ml cumate (4-isopropylbenzoic acid) was added the next day and the cells were harvested after another 48h.

### Western blotting

SDS-polyacrylamide gel electrophoresis and immunoblot analysis were performed using protein samples harvested with RIPA buffer (50 mM Tris/HCl pH 8.0, 0.1% SDS, 150 mM NaCl, 1% IGEPAL, 0.5% deoxycholate). For protein quantification, the Pierce Detergent Compatible Bradford Assay Reagent (Thermo Fisher Scientific) was used. Total protein loading on SDS-polyacrylamide gels was determined using TCE-staining [34]. Monoclonal mouse anti-FLAG primary antibody (F3165, 1:3000 dilution, Sigma Aldrich) and polyclonal goat anti-mouse secondary antibody (115-035-003, 1:3000 dilution, Jackson ImmunoResearch) was used to detect FLAG-tagged proteins. Detection was performed with ECL Prime Western Blotting Detection Reagent (Amersham) and the Vilber Fusion FX6 Edge imaging system (Vilber Lourmat).

### RNA extraction

For RNA extraction, the cells were harvested with 1 ml in-house prepared TRI reagent per 6 well [35] and RNA was isolated according to standard protocols. 150 μl 1-Bromo-3- chloropropane (Sigma-Aldrich) was used to induce phase separation the washed RNA pellet was dissolved in 20 μl RNase-free water by incubating for 10 min on a shaking 65 °C heat block.

### Quantitative reverse transcriptase (RT)-PCR

Reverse transcription was performed with 3 µg of total RNA in a 20 µl reaction volume with 10 µM VNN-(dT)20 primer and the GoScript Reverse Transcriptase (Promega). Probe-based multiplex quantitative RT-PCRs were performed with the PrimeTime Gene Expression Master Mix (IDT), 2% of cDNA per reaction, and the CFX96 Touch Real-Time PCR Detection System (Bio-Rad). PrimeTime qPCR Assays containing primers and probes were purchased from IDT (B2M = Hs.PT.58v.18759587, ZFAS1 = Hs.PT.58.25163607, GAS5 = Hs.PT.58.24767969) and used at 1x final concentration according to the manufacturer’s instruction. Each biological replicate was repeated in technical triplicates and the average Ct (threshold cycle) value was measured. The housekeeping gene B2M (FAM-labelled) Ct values were subtracted from the target (ZFAS1, Cy5-labelled or GAS5, SUN-labelled) values to receive the ΔCt. To calculate the mean log2 fold changes three biologically independent experiments were used. The log2 fold changes are visualized as single data points and mean.

### Computational analyses of RNA-sequencing data

Five publicly available RNA-sequencing datasets were obtained and analyzed: 1) SMG7-KO + SMG5/SMG6-KD in HEK293 cells (E-MTAB-9330, ArrayExpress database at EMBL-EBI, [36]), 2) GFP-CoV-2-N overexpression in HEK293 cells (GSE171010, Gene Expression Omnibus (GEO), [37]), 3) CoV-2-N-2xStrep overexpression in HEK293 cells (PRJEB44716, European Nucleotide Archive, [38]), 4) CoV-2-N-2xStrep overexpression in Calu-3 cells (PRJEB45515, European Nucleotide Archive, [38]) and 5) SARS-CoV-2 infection in Calu-3 cells (GSE148729, Gene Expression Omnibus (GEO), [39]). Transcript abundance estimates were computed with Salmon (version 1.9.0) [40] with a decoy-aware transcriptome based on GRCh38 GENCODE release 42 transcript annotations [41]. After the import of transcript abundances in R (https://www.R-project.org/ version 4.3.0) using tximport [42] (version 1.28), differential gene expression analysis was performed with the DESeq2 R package [43] (version 1.40.1). Genes with less than 10 counts in half the analyzed samples were pre-filtered and discarded. The DESeq2 LFC estimates were shrunk using the apeglm R package [44] (version 1.22.1). Differential transcript expression analysis was performed using the Swish method from the fishpond R package [45] (version 2.6.2) based on 30 inferential replicate datasets drawn by Salmon using Gibbs sampling and imported via tximeta [46]. Transcripts were pre-filtered using 10 counts per transcript in at least one condition as cut-off. Significance cut-offs were log2FoldChange > 1 & padj < 0.0001 for DESeq2 DGE and log2FC > 1 & qvalue < 0.01 for Swish DTE. All plots were generated using ggplot2 [47] (version 3.4.2) or nVennR [48] (version 0.2.3).

## Results

### SARS-CoV-2 N directly interacts with the NMD factor UPF2

To recapitulate the protein-protein interactions identified in the AP-MS study described above *in vitro*, we performed GST-pulldown assays using full-length GST-tagged N (GST-N) as a bait and UPF1, UPF2 and MOV10 as preys. For this study, we used a construct of UPF1 comprising the cysteine-histidine rich (CH) domain and the helicase core, and lacking the N- and C-terminal unstructured regions. Since full-length UPF2 is proteolytically unstable, we used a truncated construct lacking 120 and 45 residues at the N- and C-termini, respectively and refer to this as UPF2 (Figure 1A). In lieu of full-length MOV10, a fusion (MOV10fus) of the unique MOV10 N-terminus (residues 1-264) with the highly homologous helicase core of UPF1 (residues 295-914) was employed. We found that the N-protein interacts with neither UPF1 nor MOV10fus but instead shows a robust interaction with UPF2 (Figure 1B). A full-length UPF1 protein (UPF1fl) expressed and purified using a baculovirus system also does not bind N *in vitro*, suggesting that there is, at best, a weak direct interaction between the N-protein and UPF1 (Supplementary Figure 1A).

**Figure 1.**
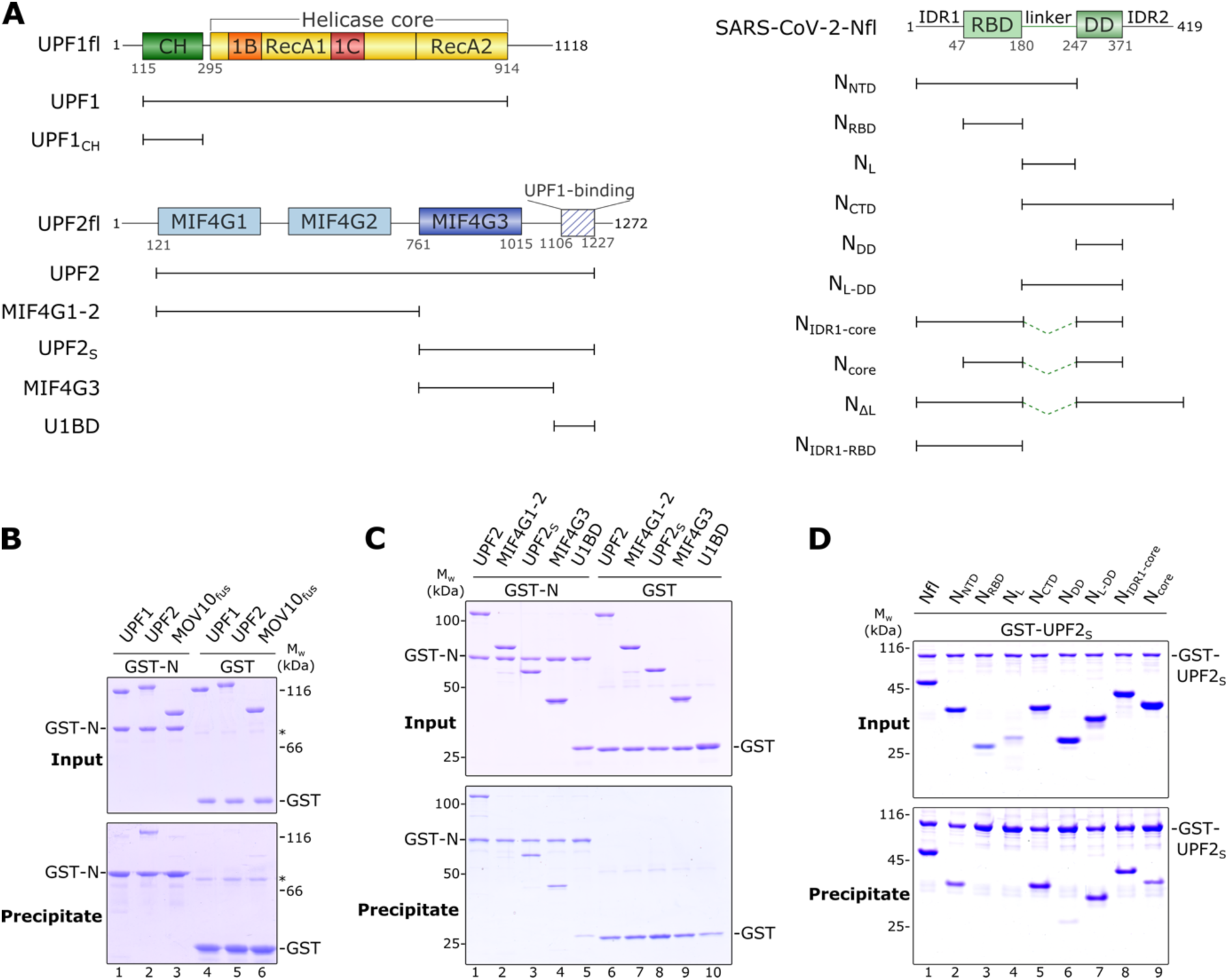
The SARS-CoV-2 Nucleocapsid (N) protein directly interacts with the core NMD factor UPF2. **A)** Schematic representation of the domain organization of human UPF1, UPF2 and SARS-CoV-2 N, and the constructs used in this study. Structured domains are shown as filled rectangles while intrinsically disordered regions (IDRs) are indicated by lines. **B)** GST-pulldown assay of UPF1, UPF2 and MOV10fus (a fusion of the N-terminus of MOV10 with the helicase core of UPF1) with GST-N as a bait. GST was used as a negative control in all such assays. The asterisk (*) indicates a contaminant. The top and bottom panels depict the inputs and precipitates, respectively, in this and all other GST-pulldown experiments. GST- N binds UPF2 but not UPF1 or MOV10fus. **C)** GST-pulldown assay to determine the domain of UPF2 that interacts with N. The MIF43G domain is the primary binding site for N, with additional weak interactions mediated by the UPF1-binding domain, U1BD **D)** GST-pulldown assay to identify the UPF2-binding region of N. Two IDRs (IDR1 and the inter-domain linker) and the dimerization domain (DD) of N make up a composite binding site for UPF2. No single site on N can mediate a strong interaction with UPF2 but a combination of any two binding sites restores binding comparable to that of full-length N (see also Supplementary Figure 1B). A complete gel including negative controls with GST is shown in Supplementary Figure 1C.

UPF2 is a multidomain protein with three ***m***iddle of e***IF4G*** (MIF4G) domains, followed by a stretch of low-complexity sequence that is natively unstructured (Figure 1A) [49]. This unstructured C-terminal region binds UPF1 and is therefore referred to as the UPF1-binding domain (U1BD) [30, 50]. Generally, MIF4G domains are well-known protein-protein interaction platforms. Although no binding partners for the MIF4G1 and 2 domains of UPF2 have been identified till date, the third mIF4G domain interacts with various RNA-processing factors such as the NMD factors UPF3 and SMG1, the double-stranded RNA binding protein Stau1 and the eukaryotic release factor eRF3 [49, 51–54]. In order to identify the binding site for N on UPF2, we made use of a series of UPF2 constructs, systematically encompassing or lacking each domain, and tested the interaction of these truncated proteins with full-length N by GST- pulldown assays. GST-N showed robust binding to all UPF2 constructs that contain the MIF4G3 domain (UPF2, UPF2S and UPF2-MIF4G3) but showed no binding to the MIF4G1-2 domains alone (Figure 1C, lanes 1-4). GST-N also bound UPF2-U1BD, albeit weaker than to constructs including the MIF4G3 domain (Figure 1C, compare lane 5 with lanes 3 and 4).

We next proceeded to map the UPF2-binding site on N using a similar approach as that described above. The N-protein has two structured domains: an RNA-binding domain (RBD) and a dimerization domain (DD) that are connected by a linker of low complexity sequence. The RBD and DD are also flanked at the N- and C-terminus, respectively, by intrinsically disordered regions (IDRs) (Figure 1A). We generated a series of truncation constructs of N that either comprise an individual domain or a domain together with its flanking IDRs (Figure 1A), and tested the interaction of these constructs with GST-UPF2S in a GST-pulldown assay (Figure 1D). As expected, full-length N (Nfl) showed strong binding to UPF2. Variants of N comprising the structured domains alone (RBD and DD) displayed no appreciable binding to UPF2: while no interaction was observed with the RBD, the DD showed a weak binding to UPF2 (Figure 1D, lanes 3 and 6). However, addition of the flexible linker and the flanking IDR to the structured domains (IDR1 in case of NRBD and IDR2 for NDD) restored the interaction with UPF2 in the resultant proteins (NNTD and NCTD, Figure 1D, lanes 2 and 5). Although neither the linker nor the DD of N show strong binding to UPF2 on their own, a variant comprising both segments (NL-DD) can mediate a robust interaction (Figure 1D, compare lanes 4, 6 and 7). As the extent of binding of the NCTD and NL-DD variants to UPF2 are very similar, we deduce that IDR2 does not contribute significantly to the N-UPF2 interaction. Overall, it appears that the N-protein engages UPF2 in a multipartite interaction, with binding sites located within IDR1, the linker and DD but not in the RBD. We hypothesize that no individual site is sufficient for binding but a combination of two sites mediates an interaction with UPF2.

To test this hypothesis and to discern the contributions of IDR1 and the linker of N in binding to UPF2, we generated two variants of the N-protein where the 67-residue long linker connecting the RBD and DD was replaced by a short 12-residue stretch consisting of glycine, serine and alanine residues (GSA-linker). The length and position of the linker was based on available high-resolution X-ray crystal structures of the RBD and DD of N as well as a multiple sequence alignment of the nucleocapsid proteins from four betacoronaviruses (Supplementary figure 1D). The first of these N variants included IDR1 in addition to the RBD and DD (referred to as NIDR1-core) while the second variant lacked IDR1 and comprised only the structured core domains (Ncore, Figure 1A). As the inter-domain linker is prone to proteolysis [55], the modified N-proteins lacking the long linker have a higher proteolytic stability than full-length N. GST-pulldown assays with these N variants showed that replacing the long linker with the short GSA-linker in NIDR1-core did not compromise its binding to UPF2 (Figure 1A, lane 8). However, deletion of IDR1 in addition to removal of the long linker led to reduced binding (Figure 1A, lane 9), supporting our previous observations that a combination of any two binding sites on the N-protein is sufficient for mediating a strong interaction with UPF2. Correspondingly, an N variant containing only the IDR1 binding site (NIDR1-RBD) shows weak binding to UPF2 (Supplementary figure 1B).

We used isothermal titration calorimetry (ITC) for a quantitative assessment of the interaction of UPF2 with N. As the full-length proteins are not stable over the duration of the measurements, we used the NIDR1-core and UPF2S proteins for ITC experiments. Furthermore, Nfl has a strong tendency to oligomerize and form unspecific higher-order assemblies in solution, precluding its use in ITC. Deletion of the long linker yielded a stable dimer of the resultant N-protein, NIDR1-core (Supplementary figure 2A). The NIDR1-core protein binds UPF2S with an affinity of 2.9 μM (Figure 2A, left panel). Surprisingly, we did not detect any binding of Ncore (lacking IDR1 and the linker) to UPF2S in ITC despite observing a weak interaction in GST-pulldown assays (Figure 2A, right panel). Our results suggest that the three binding sites, IDR1, linker and DD, form a composite binding interface and at least two of the three sites are essential for N to stably engage UPF2.

**Figure 2.**
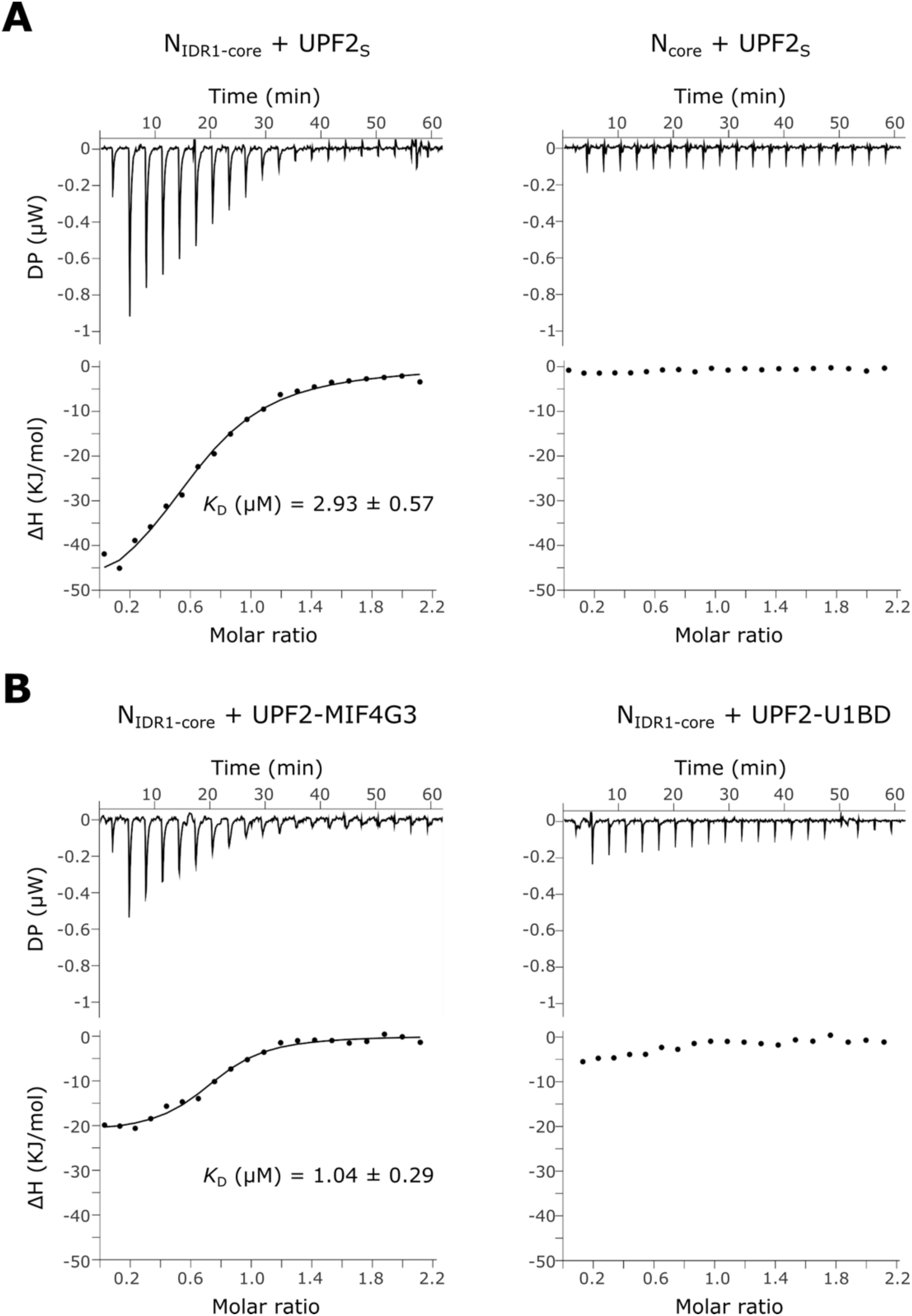
The MIF4G3 domain of UPF2 stably associates with an N variant comprising two of the three identified binding motifs. **A)** Isothermal titration calorimetry (ITC) experiments of UPF2S with NIDR1-core (left panel) and Ncore (right panel). 200 μM of UPF2S was injected into a solution of 20 μM N-protein in consecutive injections of 2 μL each. The dissociation constant calculated from the binding isotherm is shown wherever applicable. Deletion of IDR1 in addition to the inter-domain linker abrogates binding of N to UPF2. **B)** ITC experiments of NIDR1-core with the MIF4G3 domain (left panel) and the U1BD (right panel) of UPF2. The experiment was conducted as described above. Removal of the U1BD does not impact binding of UPF2 to N (compare with left panel of Figure 2A), consistent with the observation that the U1BD shows no appreciable affinity for N. All N variants are considered as a dimer in this and subsequent experiments (see Supplementary figure 2A).

Our observations with the different N constructs in ITC prompted us to further investigate the binding of N to the different UPF2 domains using ITC. GST-pulldown assays described above indicate that the MIF4G3 domain of UPF2 is the primary binding site for the N-protein while additional weak interactions are mediated by the U1BD. Using ITC, we found that the UPF2- MIF4G3 domain alone bound the N-protein with an affinity comparable to that of UPF2S (Figure 2B, left panel). However, no binding of the U1BD to the NIDR1-core was detected (Figure 2B, right panel). Conversely, addition of UPF1CH (which binds the U1BD) to UPF2S did not influence the binding affinity of N for UPF2 (Supplementary figure 2B). Taken together, our results indicate that the interaction between N and UPF2 is driven predominantly by the MIF4G3 domain of UPF2.

### SARS-CoV-2 N engages the RNA helicase UPF1 via UPF2 and modulates its catalytic activity

As the contribution of the U1BD to the N-UPF2 interaction appeared to be negligible, we proposed that UPF2 simultaneously engages UPF1 and N and thereby act as an adaptor to bridge the two proteins. To test this, we performed an analytical size-exclusion chromatography (SEC) assay using the NIDR1-core protein, UPF2S and UPF1. As predicted, UPF2 can simultaneously bind UPF1 and the N-protein to form a ternary complex, indicated by its lower retention volume compared to that of the UPF1-UPF2 complex (Figure 3A, top panel, compare black and blue traces). However, despite addition of an equi-molar amount of N, the association of N with the UPF1-UPF2 complex is sub-stoichiometric, as indicated by the weaker intensity of N on SDS-PAGE and the second peak (peak 2) containing free N. Furthermore, it appears that a small fraction of UPF1 is also released from the complex as some UPF1 protein migrates at a higher retention volume, independent of the ternary complex (Figure 3A, peak 2 and lanes 4-5 in top and bottom panels, respectively). It appears that although N can associate with UPF1 via UPF2, binding of N to UPF2 likely perturbs the UPF1- UPF2 interaction. Since the ATPase and nucleic acid-unwinding activities of UPF1 are stimulated upon binding UPF2, we hypothesized that the presence of N would impact UPF1 activity. To this end, we set up a nucleic acid-unwinding assay based on previous reports by Fritz and co-workers [56]. Briefly, a 93 nucleotide (nt) long RNA strand was annealed at its 3’- end to an 18-nt complementary DNA strand labelled with a fluorophore at its 5’-end to create a fluorescence-labeled RNA:DNA duplex. Translocation of UPF1 on the RNA strand in an ATP-dependent manner resulted in unwinding of the duplex and release of the fluorescent DNA, which was efficiently captured and quenched by a Black hole quencher (BHQ1)-labelled trap strand complementary to the DNA. The resultant decrease in fluorescence was quantified as a measure of UPF1 activity (Figure 3B, yellow trace). Addition of UPF2S to the reaction led to a rapid decrease in fluorescence, consistent with its role in stimulating UPF1 catalytic activity (Figure 3B, compare yellow and blue traces). Addition of an equi-molar amount of N to the mixture of UPF1 and UPF2S led to a significant reduction of UPF1 unwinding activity. The extent of inhibition by N was proportionate to the amount of N-protein added to the reaction (Figure 3B, green traces). Surprisingly, only 1/4^th^ the molar amount of N with respect to UPF1 (molar ratio of N:UPF1 is 1:4) was sufficient to mediate a small reduction in unwinding activity (Figure 3B, pale green trace). RNA-dependent ATPase assays of a mixture of UPF1- UPF2S in the presence of varying amounts of N showed a similar trend, with a small reduction in activity upon addition of limiting (0.25X) amounts of N-protein and robust inhibition in presence of excess (2X) N (Figure 3C). Taken together, our observations show that interaction of N with UPF2 impairs the ability of UPF2 to stimulate UPF1 catalytic activity. At first glance, this appears incongruous as N and UPF1 bind two distinct domains of UPF2, the MIF4G3 and U1BD, respectively, and activation of UPF1 by UPF2 is a consequence of a conformational change induced upon interaction. However, our ATPase assays show that the U1BD of UPF2 alone cannot activate UPF1. The MIF4G3 domain is essential for robust activation of UPF1, although it does not exert this effect alone either (Figure 3D). We suggest that binding of the U1BD of UPF2 to UPF1 positions the UPF2-MIF4G3 domain close to the helicase core of UPF1, which has a positive impact on UPF1 catalytic activity. Binding of N to the UPF2- MIF4G3 domain interferes with this effect and prevents complete activation of UPF1 by UPF2.

**Figure 3.**
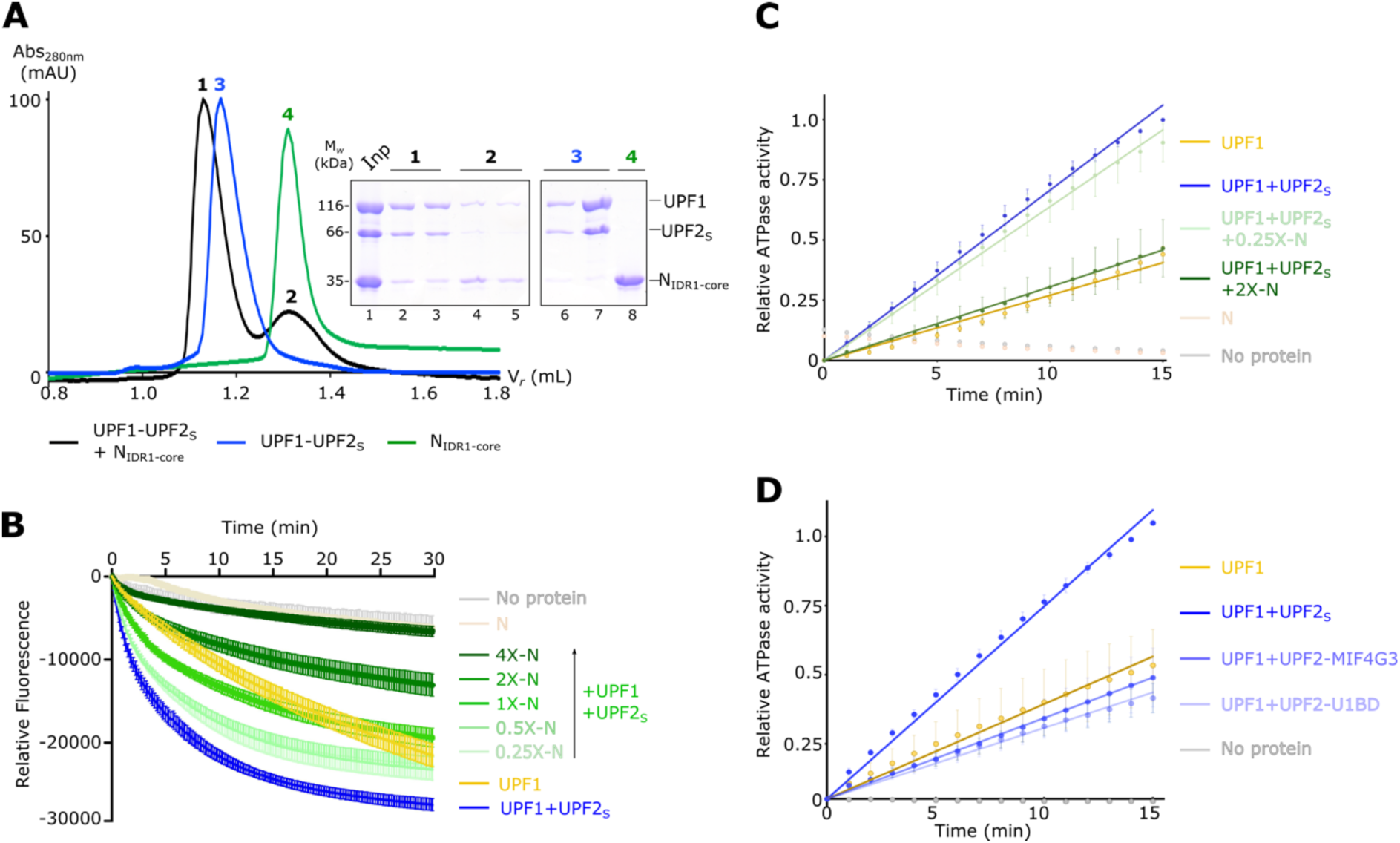
Indirect association of N with UPF1 inhibits its catalytic activity. **A)** Analytical size-exclusion chromatography (SEC) depicting formation of a ternary complex of UPF1, UPF2S and NIDR1-core. The terms Abs and V*r* refer to absorbance (at 280 nm or 260 nm, as indicated in the figure) and retention volume, respectively, in this and all other figures. The exclusion volume of the column is 0.8 mL. The corresponding SDS-PAGE analysis of the peak fractions, visualized by Coomassie-staining, is shown below. Interaction of N with the UPF1- UPF2 complex is sub-stoichiometric and leads to release of a small amount of UPF1 from the complex. **B)** Fluorescence-based nucleic acid-unwinding activity measurements of a mixture of UPF1 and UPF2S in presence of increasing concentrations of N (0.25- to 4-fold excess over UPF1). Measurements were taken at 10 s intervals for a total duration of 30 minutes. The data and the associated error bars represent the mean and standard deviation of 3 independent experiments. Technical duplicates were performed for each experiment. Controls without any protein and with N alone were included to monitor the stability of the RNA:DNA hybrid over the course of the experiment. The unwinding activity of UPF1 alone is shown for comparison. A low amount of N is sufficient to initiate inhibition of UPF1 unwinding activity, which is completely blocked in presence of high amounts of N. **C)** RNA-dependent ATPase activity of a mixture of UPF1 and UPF2S in the presence of low (0.25-fold of UPF1) and high (2-fold excess over UPF1) concentrations of N. An enzyme-coupled phosphate detection assay was used to measure ATPase activity. The data and the associated error bars represent the mean and standard deviation of 3 independent experiments. Technical duplicates were performed for each experiment. As with the unwinding activity measurements, a low amount of N slightly inhibits UPF1 ATPase activity while a higher amount shows more robust inhibition. **D)** RNA-dependent ATPase activity of UPF1 in presence of different variants of UPF2 (added in 2-fold excess over UPF1). Although a UPF2 variant comprising the MIF4G3 and the U1BD domains strongly stimulates the ATPase activity of UPF1 (UPF2S, dark blue trace), these domains are not capable of activating UPF1 on their own. Experimental setup and data presentation are as described above in (C).

### SARS-CoV-2 N displaces UPF1 from RNA at higher concentrations and indirectly inhibits its catalytic activity

The observations described above suggest that the influence of N on the catalytic activity of UPF1 can only be exerted through UPF2. However, activity assays of UPF1 alone showed that when present in excess, N can directly inhibit both unwinding as well as the ATPase activities of UPF1 (Figures 4A and 4B). Unlike with UPF1-UPF2, low amounts of N do not modulate the catalytic activity of UPF1 alone. Although N specifically binds sequence and structural elements at the 5’-UTR of the SARS-CoV-2 genomic RNA [57, 58], *in vitro* it can bind a generic homopolymeric sequence (such as poly-U) and mediate liquid-liquid phase separation [59]. Given that UPF1 also binds RNA without any sequence specificity, we asked if N and UPF1 compete for binding RNA and if this influences the catalytic activity of UPF1. We first used fluorescence anisotropy to determine the binding affinity of N for a 12-mer poly- U RNA (U12) labelled with 6-FAM at its 5’-end. The binding affinity of Nfl as well as NýL for RNA is comparable to that previously determined for UPF1 (*K*D of ∼ 50 nM for UPF1-RNA vs 82 nM for Nfl-RNA and 45 nM for NýL-RNA, Supplementary figure 4A). Analytical SEC assays of a mixture of UPF1, a 45-mer poly-U RNA (U45) and 2-fold molar excess of NýL showed that N and UPF1 can co-occupy the U45 RNA, as indicated by a single peak (peak 1) containing both proteins as well as the RNA (Figure 4C, peak 1 and bottom panel lanes 1 and 2). The retention volume of peak 1 is lower than that of the peaks corresponding to the N-RNA and UPF1-RNA complexes, respectively, confirming that it represents a protein-RNA complex comprising both N and UPF1. However, the amount of N in peak 1 is higher than that of UPF1, based on the intensities of the bands on the Coomassie-stained gel. Additionally, we observed an additional peak (peak 3) corresponding to free UPF1, which has a lower absorbance at 260 nm than at 280 nm and correspondingly, a lower amount of RNA (Figure 4C, peak 3 and lane 4 of top and bottom panels, respectively). The amount of free UPF1 obtained depends on the fold- excess of N-protein added, where excess N stimulates a greater release of UPF1 from RNA (Supplementary figure 4B). Our results suggest that although N binds RNA with an affinity comparable to UPF1 and there is no direct competition for RNA binding, excess N can impede RNA-binding by UPF1, rendering it incapable of translocating on RNA and mediating its ATPase and unwinding activities.

**Figure 4.**
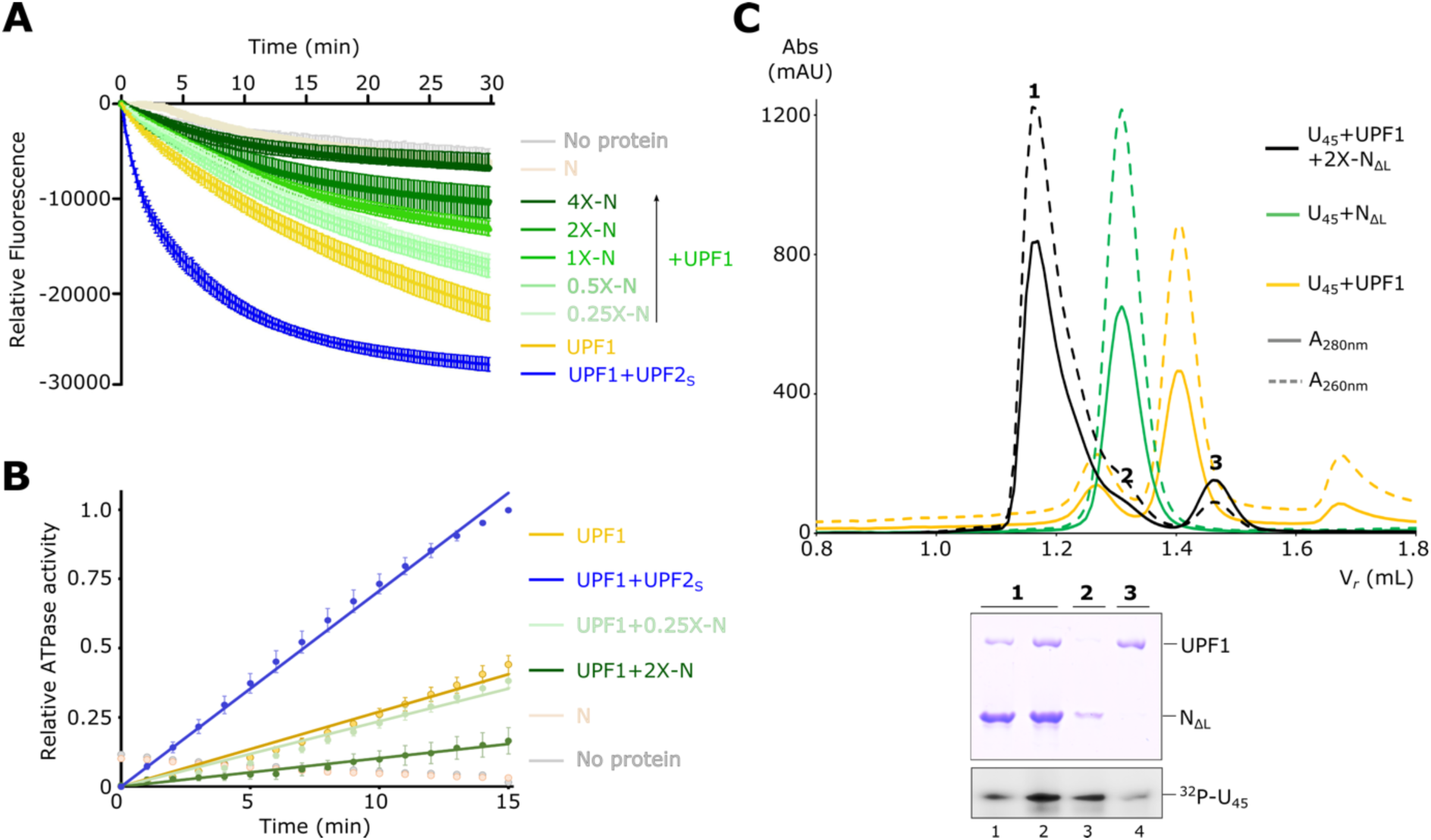
High concentrations of N impede UPF1 catalytic activity by displacing it from RNA. **A)** Nucleic acid-unwinding activity of UPF1 in presence of increasing concentrations of N (0.25- to 4-fold excess over UPF1). The experiment, including controls, was conducted as described in Figure 3B. The unwinding activity of a mixture of UPF1 and UPF2S is shown for comparison. In absence of UPF2, N inhibits UPF1 activity only at high concentrations. **B)** RNA-dependent ATPase activity of UPF1 in presence of low (0.25-fold) and high (2-fold excess) concentrations of N. As observed with unwinding activity measurements, inhibition is only achieved upon addition of high amounts of N. **C)** Analytical SEC experiments of UPF1 and U45 RNA in the presence of 2-fold excess of Nι1L. An overlay of the chromatogram of the U45-UPF1- N mixture (black) with those of U45-N (green) and U45-UPF1 (yellow) are shown in the top panel. The bottom panels show the corresponding SDS- and urea-PAGE analyses of peak fractions of the U45-UPF1-N chromatogram. Proteins were visualized by Coomassie staining and RNA was detected by radiolabeling with ^32^P followed by phosphorimaging. Addition of an excess of N to a UPF1-RNA mixture leads to higher occupancy of N on RNA (peaks 1 and 2) and a concomitant release of UPF1 from RNA (peak 3).

### Impact of SARS-CoV-2 N on host cell NMD targets

Our biochemical and biophysical results suggest that the SARS-CoV-2 N-protein can potentially interfere with the proper cellular function of UPF1 in NMD. To test this hypothesis, we generated stably transfected Flp-In-T-REx-293 cells, which express N-terminally FLAG-tagged proteins in a cumate-inducible manner. We used emGFP as negative control and a previously reported dominant negative UPF1 mutant (R843C) as positive control [60]. In addition, we tested two variants of N proteins (bat = SARS-like bat-SL-CoVZC45; human = SARS-CoV-2) which exhibit 94% protein identity. After 48h of induction, western blot experiments confirmed the expression of all proteins, with CoV-2-N (human) and UPF1- R843C exhibiting the strongest expression levels (Figure 5B and Supplemental figure 5A). Despite the comparable expression levels, only the UPF1-R843C expression resulted in a modest upregulation of two known NMD targets (ZFAS1 and GAS5) in qPCR experiments [61], whereas both N proteins showed no effect (Figure 5C). Both ZFAS1 and GAS5 are snoRNA host genes, which are degraded by NMD, however many other classes of NMD targets exist in human cells. To analyze the transcriptome-wide impact of N overexpression on NMD activity, we analyzed publicly available RNA-Seq datasets [37, 38]. Since NMD inhibition is expected to lead to the accumulation of NMD-targeted transcripts, we compared the significantly upregulated genes between three different N-overexpressing conditions (two in HEK293, one in Calu-3 cells) and in relation to a strong NMD inhibition (SMG7 knockout + SMG5/SMG6 knockdown, [36]) Overall, the number of overlapping genes between the different studies is low and only 2 genes are consistently found as significantly upregulated in all conditions (IFIH1 and IFI44L; Figure 5D). Both genes are implicated in the defense response to viruses (gene ontology term GO:0051607) and have at least one NMD-annotated transcript in the GENCODE database. When investigating five other known commonly upregulated NMD targets, we do not find compelling evidence that overexpressed N protein acts as a broad NMD inhibitor (Figure 5E). Furthermore, no significant upregulation of those NMD targets could be observed in RNA-Seq data from Calu-3 cells infected with SARS-CoV- 2 for different timepoints (4 to 12 hours, Figure 5E, [39]). Next, we analyzed the differential transcript expression (DTE) and asked whether NMD-annotated transcripts are preferentially upregulated. Whereas this is very apparent for the strongly NMD-inhibited SMG7-KO+SMG5- KD conditions, N overexpression did not result in substantial NMD-isoform accumulation (Figure 5F and Supplemental figure 5B). Taken together, our analyses of various overexpression studies could not identify a widespread NMD-inhibitory role of SARS-CoV-2 N in cultured human cell lines.

**Figure 5.**
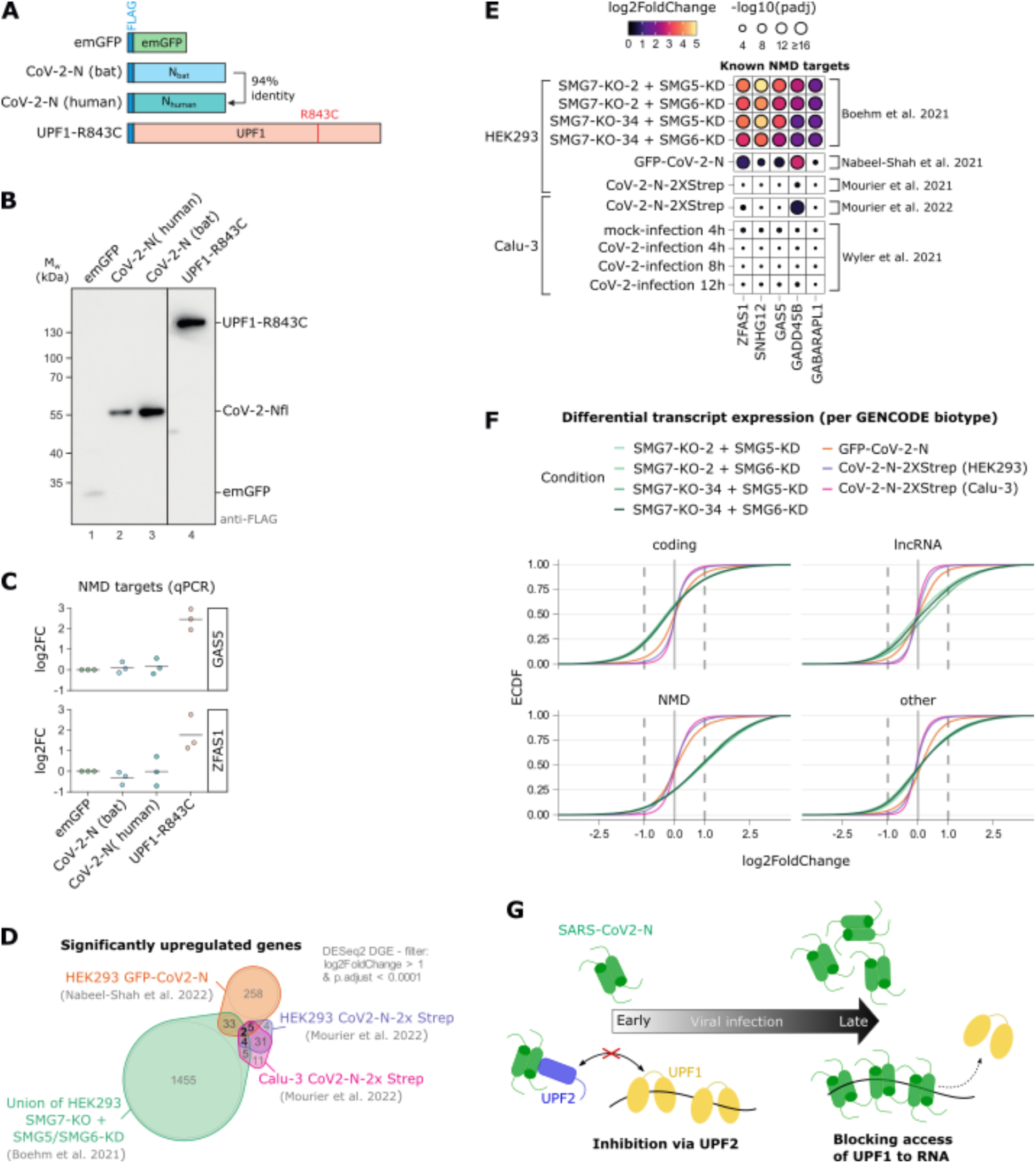
NMD is not substantially inhibited by N overexpression in human cell lines. **A)** Schematic representation of the overexpressed FLAG-tagged constructs. **B)** Western blot analysis of FLAG-tagged protein expression in HEK293 cells (n=3). **C)** Probe-based quantitative RT-PCR analysis of GAS5 and ZFAS1 expression levels in the respective overexpression condition normalized to the B2M reference. FLAG-emGFP serves as negative and UPF1-R843C as positive control. Data points and means are plotted as log2 fold change (log2FC) (n=3). **D)** Overlap of significantly upregulated genes (cut-offs are indicated) in RNA-Seq data from three different CoV-2-N overexpression datasets and the union of four NMD-factor knockout/knockdown combinations. **E)** Heatmap of DESeq2-derived log2 fold changes of five selected known NMD target genes in various RNA-Seq datasets. The size of points corresponds to the statistical significance. The CoV2-N-2XStrep dataset is obtained from the preprint https://www.medrxiv.org/content/10.1101/2021.05.06.21256706v2 (Mourier et al., 2021) **F)** Cumulative distribution function plots of differential transcript expression determined by Swish and stratified by GENCODE biotype (coding = protein-coding). **G)** A mechanistic model for inhibition of UPF1 catalytic activity on SARS-CoV2-N RNA by low and high amounts of N-protein in host cells. The amount of N in host cells continuously increases with progression of infection.

## Discussion

The influence of NMD factors on the replication of positive-sense RNA viruses has been previously investigated in detail. Depletion of NMD factors leads to increase in the efficiency of replication and propagation of the Semliki Forest virus (SFV) [62]. The viral genomic RNA as well as sub-genomic mRNA of MHV was shown to be recognized by the NMD pathway and consequently, stabilized upon inhibition of NMD [18]. As with SFV, depletion of NMD factors leads to an enhancement in MHV replication. It follows that these viruses must therefore develop mechanisms to inhibit NMD in order to facilitate their replication and propagation. The capsid protein of SFV and the nucleocapsid protein (N) of MHV were shown to inhibit the NMD pathway, suggesting a direct intervention of the viral proteins in this decay mechanism [18, 63]. In this study, we show that the nucleocapsid protein of the SARS-CoV-2 virus directly interacts with the core NMD factor UPF2. We find that the N-protein can associate with UPF1 via interactions with UPF2, accounting for the observed interaction between the two proteins in a high-throughput interactome study.

The N-protein uses a combination of IDRs and a structured domain to interact with the MIF4G3 domain of UPF2. IDRs are often involved in mediating protein-protein interactions due to their intrinsic flexibility and their ability to adopt an ensemble of different conformations. They can adopt a distinct fold upon binding a structured domain or can engage in fuzzy interactions where the conformation of the IDR in the complex remains heterogenous. IDRs can also interact with unstructured proteins/region to form complexes of diverse stabilities. The inter- domain linker and IDR1 of N together with its C-terminal dimerization domain form a composite binding surface for UPF2. Interestingly, any two of the three interfaces of N are sufficient to mediate a robust interaction with UPF2. The inter-domain linker alone probably has the least contribution to binding as its removal does not significantly reduce binding to UPF2 (Figure 1D). The linker was shown to play an important role in the higher order oligomerization of N, which is critical for packaging the viral genome and stabilizing the genomic RNA. It is therefore plausible that the additional interactions mediated by the linker with host cell proteins such as UPF2 are weaker than those forged by IDR1 and the DD as it is already involved in *cis*- intermolecular interactions.

UPF2 plays multiple roles in NMD. It directly binds UPF1 and is believed to engage the helicase in a large assembly referred to as the decay-inducing complex (DECID) that involves the exon junction complex (EJC) and the eukaryotic release factors (eRFs) [54, 64–66]. It was also thought to facilitate phosphorylation of UPF1 which is crucial for NMD in metazoans. Additionally, binding of UPF2 to UPF1 mediates its release from RNA and stimulates its catalytic activity, which is essential for NMD [31]. The modular domain arrangement of UPF2 enables these functions as it uses distinct domains to associate with UPF3-EJC on one hand, and with UPF1 on the other. Binding of UPF3 to the UPF2-MIF4G3 domain does not impact the association of UPF1 with UPF2 [50]. Similarly, concomitant binding of UPF2 to Stau1 and UPF1 in the SMD pathway allows for activation of UPF1 within the ternary complex [51]. In comparison to these assemblies, the interaction of N to UPF2 is rather unusual. The N-protein binds the MIF4G3 domain of UPF2 and can form a ternary complex, albeit sub-stoichiometric, with UPF1 and UPF2. However, instead of activation of UPF1 as is typically observed in the other ternary complexes mentioned above (UPF1-UPF2-UPF3 and UPF1-UPF2-Stau1), we observe a strong inhibition of UPF1 catalytic activity. We speculate that the MIF4G3 surface that binds N is distinct from the one that binds UPF3 and Stau1. Fittingly, our preliminary data show that UPF2 can simultaneously bind N and UPF3 in analytical SEC (Supplementary figure 3). Given that the UPF2-MIF4G3 domain has a strong positive influence on UPF1 catalytic activity (Figure 3D), it is possible that binding of N to it negates this effect and as a result, UPF1 is no longer activated by UPF2 even though it remains bound through the U1BD.

In addition to the effect mediated upon binding of N to UPF2, we observe a robust inhibition of UPF1 catalytic activity by N alone. Unlike in the presence of UPF2, this inhibitory effect is only observed with an excess of the N-protein. We attribute this to high amounts of N coating the RNA and blocking access of UPF1 to the RNA (see also Figure 4C). Based on our observations of the UPF2-dependent and UPF2-independent inhibition of UPF1 catalytic activity by N, we propose the following model (Figure 5G) – at early stages of viral infection, the levels of N-protein in the host cell are low. At this stage the genomic RNA needs to be efficiently replicated and the sg-mRNAs need to be produced in large amounts to generate the structural proteins necessary for viral packaging and propagation. Therefore, it is imperative that the genomic RNA as well as the sg-mRNA are protected from degradation by the NMD pathway and are adequately stabilized. We speculate that small amounts of N present at this stage binds UPF2 and prevents it from activating UPF1, thereby minimizing decay of the viral RNA. As infection progresses, so does the accumulation of N in the host cell [67]. Large amounts of N coat the viral RNA and prevent binding of UPF1 and the onset of NMD (Figure 5G). Despite the potent inhibitory effect of N observed *in vitro*, we do not observe transcriptome-wide stabilization of cellular NMD targets in mammalian cells. Also, human and bat N-proteins do not stabilize NMD targets in different cellular systems (Figure 5G). This is particularly interesting as the MHV-N protein was previously shown to stabilize a luciferase reporter that mimics an MHV sg-mRNA. Global shutdown of NMD would be fatal for host cells and therefore detrimental to viral replication and propagation. It is therefore intuitive that especially at early stages of viral infection, the N protein does not block the general decay of cellular NMD targets. As N binds structural elements in the 5’-regulatory region of the SARS- CoV-2 genomic RNA with high specificity, it is possible that only viral RNA containing these *cis*-acting elements are protected from NMD by N. The underlying mechanism of how the N- protein precisely inhibits NMD of viral RNA without interfering with host cell NMD targets remains a question for future studies.

## Data Availability

All original data will be made available upon request.

## Acknowledgements

We thank Nicole Holton for technical assistance with ITC, Matthias Ballauf for sharing his expertise in ITC data analysis, Silke Modersohn for preparing and conducting cell culture experiments, and Markus Wahl and Elena Conti for the expression plasmids of the SARS- CoV2 N-protein and full-length human UPF1, respectively. We are grateful to members of the Chakrabarti and Gehring labs for helpful discussions and for critical comments on the manuscript. This study was supported by the Deutsche Forschungsgemeinschaft (CH1245/3- 2, CH1245/5-1 and CH1245/6-1 to S.C. and GE2014/6-2 to N.H.G.) and core funding from Freie Universität Berlin. S.C. is supported by the Heisenberg Program of the DFG. The authors declare that there are no conflicts of interest.

## Author Contributions

M.M., G.X and M.B purified the proteins used in the study. M.M. performed the biochemical analyses and biophysical assays with assistance from G.X., M.B and S.C. V.B. and N.H.G. performed the cell-culture studies and the RNA-Seq analyses. M.M and S.C. designed the study and wrote the paper with input from V.B. and N.H.G.

## Supplementary Methods

### Expression and purification of UPF1fl from a baculovirus system

Hi5 cells at a density of 0.8x10^6^ cells/mL were infected with a first generation virus encoding for UPF1fl at a v/v ratio of 1:1000 (virus:cells). Cells were harvested 68 hours post-infection by centrifugation at 800xg for 10 mins. The cell pellet was rinsed with 1X phosphate-buffered saline (PBS) and resuspended in lysis buffer A (50 mM Tris-HCl pH 7.5, 500 mM NaCl, 10 mM imidazole and 10 % glycerol) supplemented with 1 mM phenylmethylsulphonyl fluoride (PMSF) and 0.5 mg of DNase I. Cells were lysed by low-pulse sonication. All subsequent steps were conducted as described for the purification of *E. coli* expressed proteins in the main text.

### SEC-MALS

To measure the absolute molecular mass of the SARS-CoV2-Nfl and NIDR1-core proteins, approximately 60 µg of protein at a concentration of 1 mg/ml was injected onto a Superdex 200 10/300 column (GE Healthcare) on an Agilent HPLC system coupled to miniDWAN TREOS (Wyatt Technology, Germany) and Refracto-Max520 (ERC) detectors. The protein was eluted using a thoroughly degassed running buffer containing 20 mM Tris-HCl, pH 8.0, 150 mM NaCl, 5% glycerol, 0.02% NaN3 and 2 mM DTT at room temperature. The instrument was calibrated using bovine serum albumin as a reference. Data recording and analysis was performed using the ASTRA 6.1.4.25 software (Wyatt Technology, Germany).

### Fluorescence anisotropy

For determination of the binding affinity of SARS-CoV-2 N for RNA, 5 nM of a 5’-end 6-FAM labeled 12-mer poly-U-RNA was mixed with increasing concentrations of Nfl and a modified stable construct of N-protein (NΔL) in a binding buffer (20 mM HEPES, pH 7.5, 100 mM NaCl, 1 mM MgCl2 and 100 µg/ml BSA) and incubated at 25 °C for 1 h in dark. The affinity of UPF1 for RNA was determined in the same setup. An amount of 40 µl of each sample was transferred to a flat-bottom black 384-well plate (PerkinElmer OptiPlate 384-F) to measure the fluorescence polarization at 25 °C with using a Spark multimode microplate reader (Tecan Lifesciences). The value obtained in absence of any protein (RNA alone) was considered as baseline and subtracted from all the other fluorescence polarization (F.P) values. Fluorescence anisotropy was calculated from fluorescence polarization using a formula [2*F. P/ (3-F. P)] and normalized against the value obtained for the highest N-concentration. The data presented are an average of at least three independent sets of experiment and were fitted to an equation representing one site-specific binding with Hill slope in GraphPad Prism 5.0. The error bars represent the standard deviation of each data point from the mean, while the error associated with the *K*D denotes standard error of mean (SEM).

### Fluorescence-based unwinding assay

The RNA substrate (5’-GGGACACAAAACAAAAGACAAAAACACAAAACAAAAGACAAAAACACAAA

ACAAAAGACAAAAAGCCAAAUUACCGUGUGCGUACAACUAGCU-3’) used in this helicase assay were prepared by in vitro transcription (IVT) from a linearised dsDNA template (CTAATACGACTCACTATAGGGACACAAAACAAAAGACAAAAACACAAAACAAAAGACAAAAACACAA

AACAAAAGACAAAAAGccaaattaccGTGTGCGTACAACTAGCT) using the oligos oAA82 and oAA83 (Supplementary Table 2). Fresh helicase substrate (RNA:DNA duplex at a 11:7 ratio) was prepared prior to every experiment by incubating the Alexa Fluor 488-labeled DNA `(Alexa488-GTGTGCGTACAACTAGCT)` and transcribed RNA with 2 mM MgOAc and 1x unwinding buffer (10 mM MES pH 6.5, 50 mM potassium acetate, 0.1 mM EDTA) at 95 °C for 3 mins and 30 sec, followed by slow-cooling the mixture down to 30 °C. For each replicate, a 40 µl reaction mixture was prepared which included 16.5 µl of reaction mixture and 2 µl of each protein of interest (final concentrations of 300 nM UPF1, 600 nM UPF2 and increasing concentrations of Nfl from 300 nM-1200 nM). The 3’-BHQ1-labeled quencher `(GTGTGCGTAC AACTAGC-BHQ1)` was added to the mixture at a final concentration of 0.56 µM (4X of labeled DNA) as a trap.

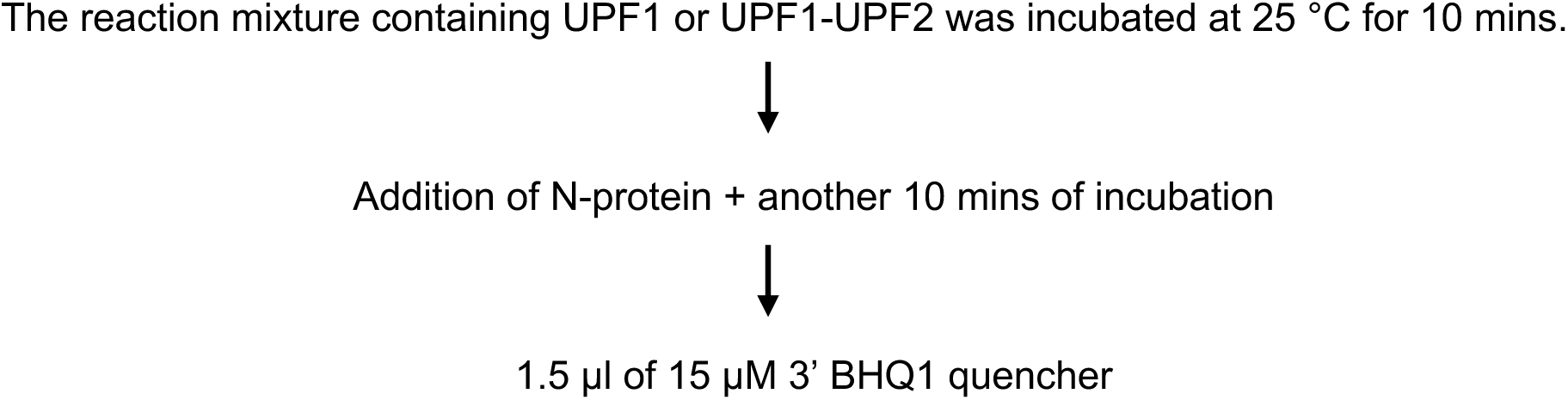

The whole mixture of 24 µl was prepared in the dark in opaque reactions tubes and subsequently transferred into a black, flat-bottomed, 384-well plate (PerkinElmer OptiPlate 384-F). 16 µl of 5 mM ATP (final concentration of 2 mM) was injected using the injector module of the plate reader. The change in fluorescence was monitored for 30 min at 30 °C. The measured fluorescence intensities were normalized to the zero time point (baseline) for each condition to obtain relative fluorescence.

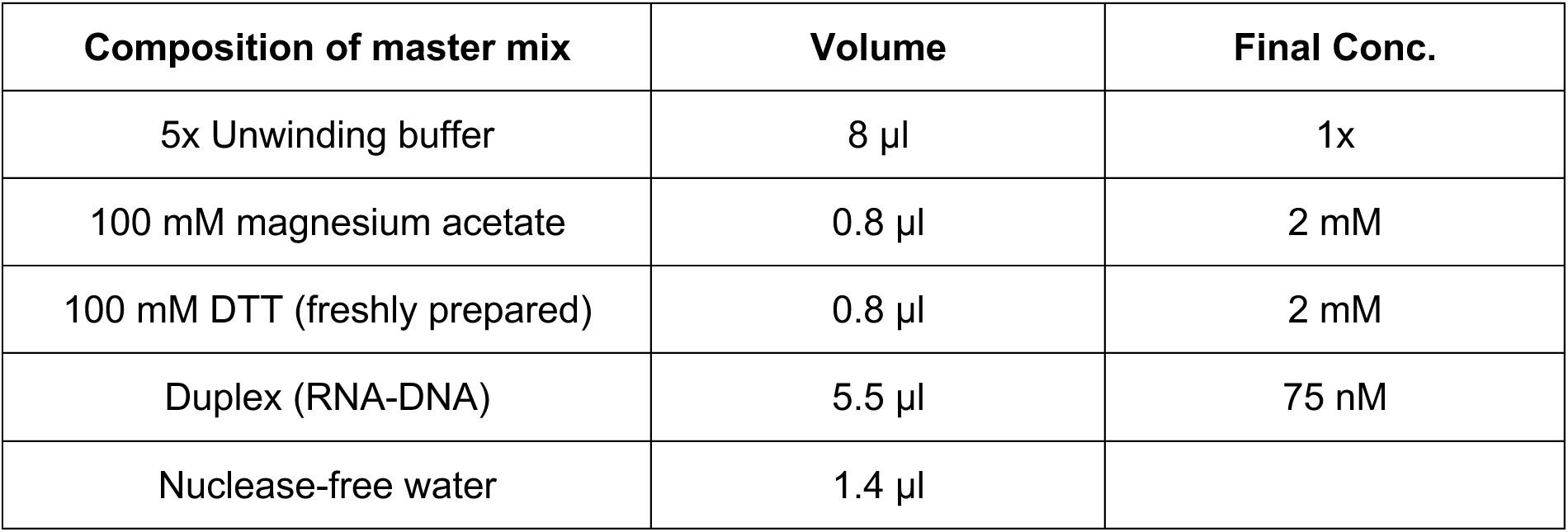

---

**Figure S1.**
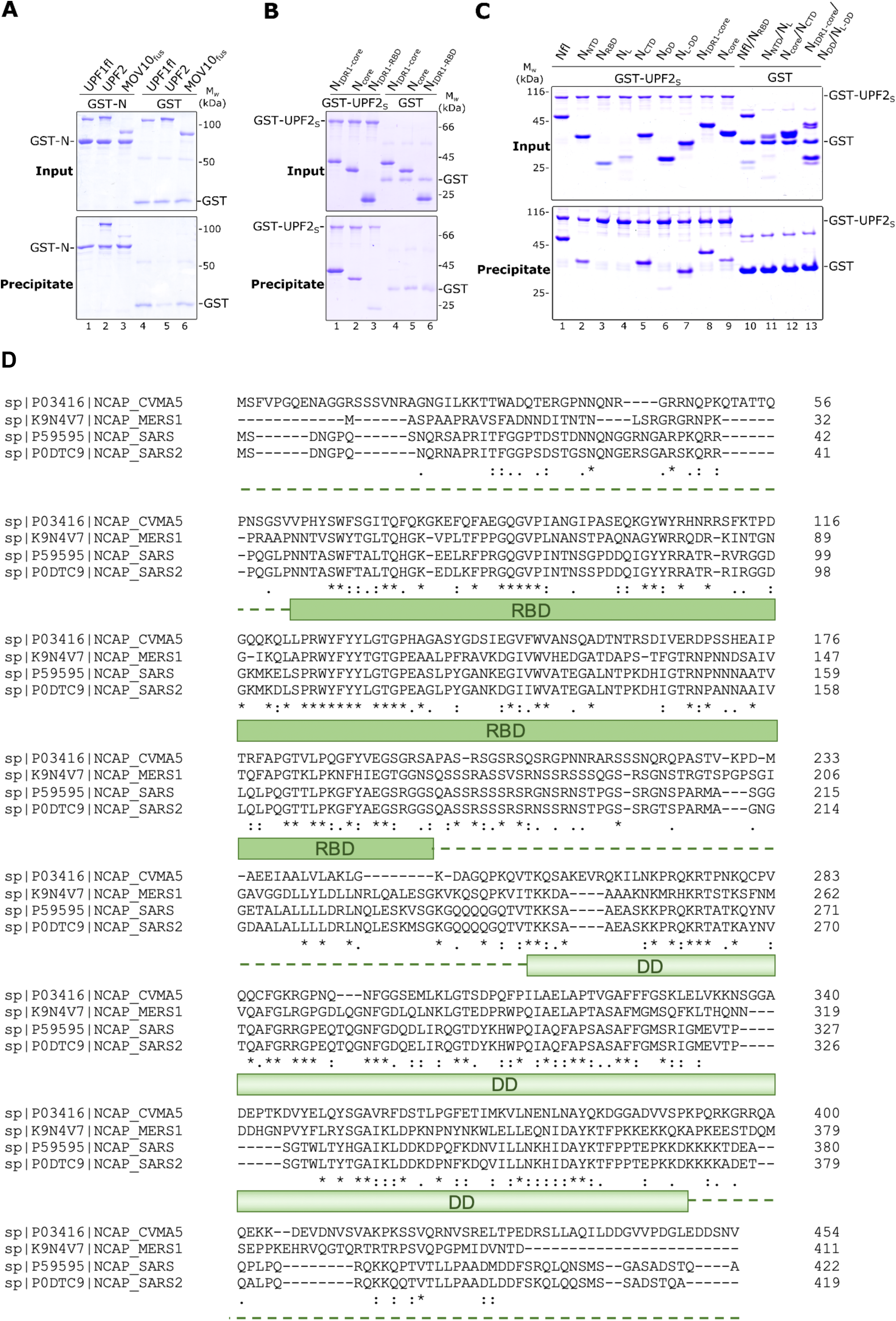
The SARS-CoV-2 Nucleocapsid (N) protein directly interacts with the core NMD factor UPF2. **A)** GST-pulldown assay of full length UPF1-(UPF1fl), UPF2, and MOV10fus (a fusion of the N-terminus of MOV10 with the helicase core of UPF1) with full-length GST-N as a bait. GST was used as a negative control in all such assays. The top and bottom panel depict inputs and precipitates, respectively, in this and all the other GST-pull down assays. GST-N binds to UPF2 but not UPF1fl or MOV10fus. **B)** GST-pulldown assay to compare binding of the IDRs and the structured domains of N to UPF2. Here GST-UPF2S is used as a bait and NIDR1-core, Ncore and NIDR1-RBD as preys. Variants of N containing only one of the three identified binding motifs (IDR1, linker and DD), such as Ncore and NIDR1-RBD, show weaker binding to UPF2 in comparison to NIDR1-core, which contains two of three binding motifs. We conclude that no single binding site of N can mediate a strong interaction with UPF2: a combination of any two binding sites are necessary for a stable N-UPF2 interaction. **C)** A complete gel corresponding to Figure 1D, including negative controls using GST as a bait. Domain organization of the N- constructs shown in Figure 1A. **D)** Multiple sequence alignment using Clustal Omega (Clustal O 1.2.4) with N-protein sequences from different coronaviruses (P03416: Murine Hepatitis Virus, K9N4V7: MERS, P59595: SARS-CoV and P0DTC9: SARS-CoV-2 strain, acquired from Uniprot: https://www.uniprot.org/). The sequence similarities across species are indicated as follows: * - fully conserved, : - strongly similar, . – weakly similar. The domain organization of SARS-CoV2 N is indicated below the alignment. Shaded boxes denote structured domains and dashed lines depict intrinsically disordered regions (IDRs) flanking the domains. Sequence conservation is highest among amino acids within the structured regions and considerably lower across the IDRs.

**Figure S2.**
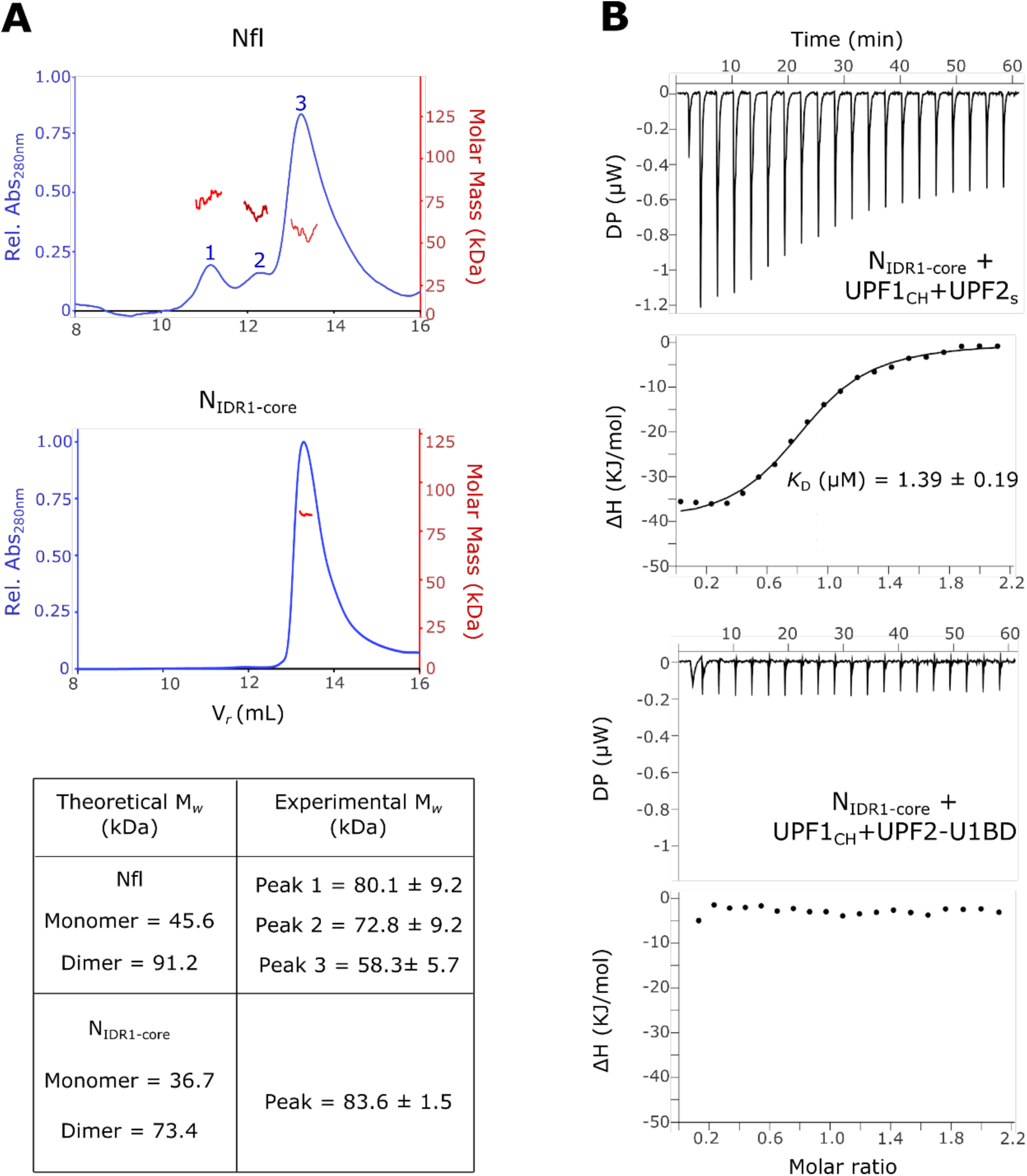
**A)** Size Exclusion Chromatography-Multi-angle Light Scattering (SEC-MALS) analysis to determine the oligomerization state of full-length N (Nfl) and NIDR1-core and the molar mass (kDa) across the observed peaks. The experimentally-obtained values are indicated in the table below, along with the theoretical molar mass (calculated using the Expasy Protparam tool) for distinct oligomerization states of each N construct. Nfl exists as a mixture of indistinct oligomers in solution whereas NIDR1-core, lacking the long inter-domain linker, forms a homogenous dimer. **B)** Isothermal titration calorimetry (ITC) experiments of NIDR1-core with complexes of UPF1CH and UPF2 (UPF2S and UPF2-U1BD, top and bottom panels, respectively) show that addition of UPF1 does not influence the binding affinity of N to UPF2 (compare *K*D derived from this experiment to that in Figure 2A).

**Figure S3.**
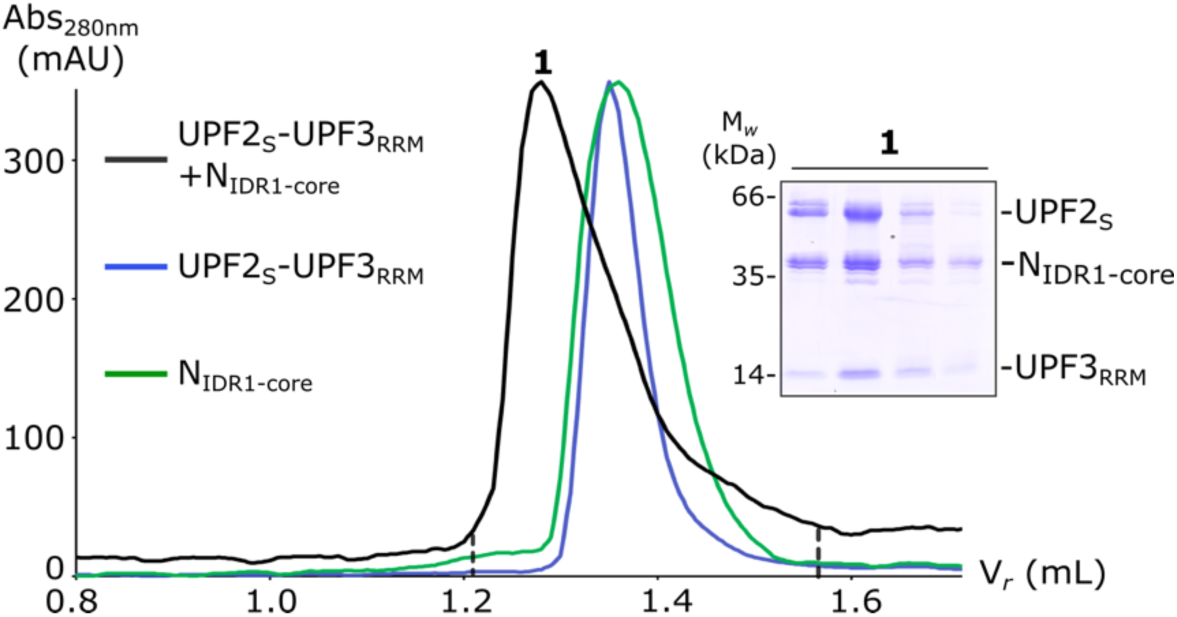
Analytical size-exclusion chromatography (SEC) shows that UPF2 binds N and UPF3 simultaneously. SDS-PAGE analysis corresponding to peak 1 of the black trace depicts formation of a ternary complex of UPF2S-UPF3RRM-NIDR1-core. The terms Abs280nm and Vr refer to absorbance at 280 nm and retention volume, respectively.

**Figure S4.**
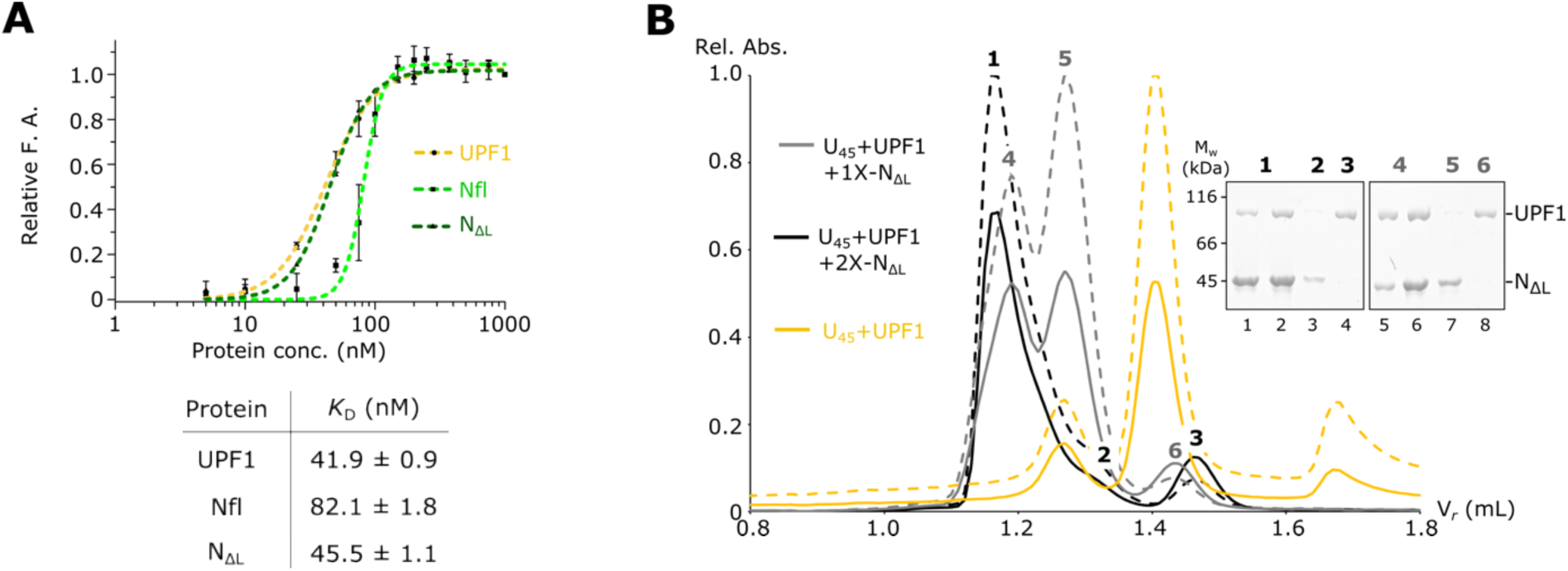
A) Fluorescence anisotropy measurements using a 5’-end 6-FAM labeled12-mer poly-U RNA (U12) shows that UPF1 (in yellow trace), Nfl (light green) and NΔL (dark green) have comparable binding affinities towards the RNA with a *K*D of ∼ 50 nM for UPF1-RNA, 82 nM for Nfl-RNA and 45 nm for NΔL-RNA. The data points and their error bars represent the mean of fluorescence anisotropy (calculated using Graph Pad Prism 5.00) and the standard deviation of independent experiments. **B)** Analytical size exclusion chromatography (SEC) and the corresponding SDS-PAGE analyses of mixtures of UPF1, a 45-mer poly-U RNA (U45) and NΔL, where UPF1 and U45 are in equi-molar amounts and NΔL is added either in equi-molar amounts to UPF1 (grey trace) and in 2-fold excess to UPF1 (black trace). Although N and UPF1 can co-occupy the RNA (lanes 1 and 5, corresponding to peaks 1 and 4), a small peak corresponding to RNA-free UPF1 is always observed (lanes 4 and 8, corresponding to peaks 3 and 6). Addition of excess N leads to an increase in free UPF1 (compare peaks 3 and 6). Solid and dashed lines refer to absorbance at 280 nm and 260 nm, respectively.

**Figure S5.**
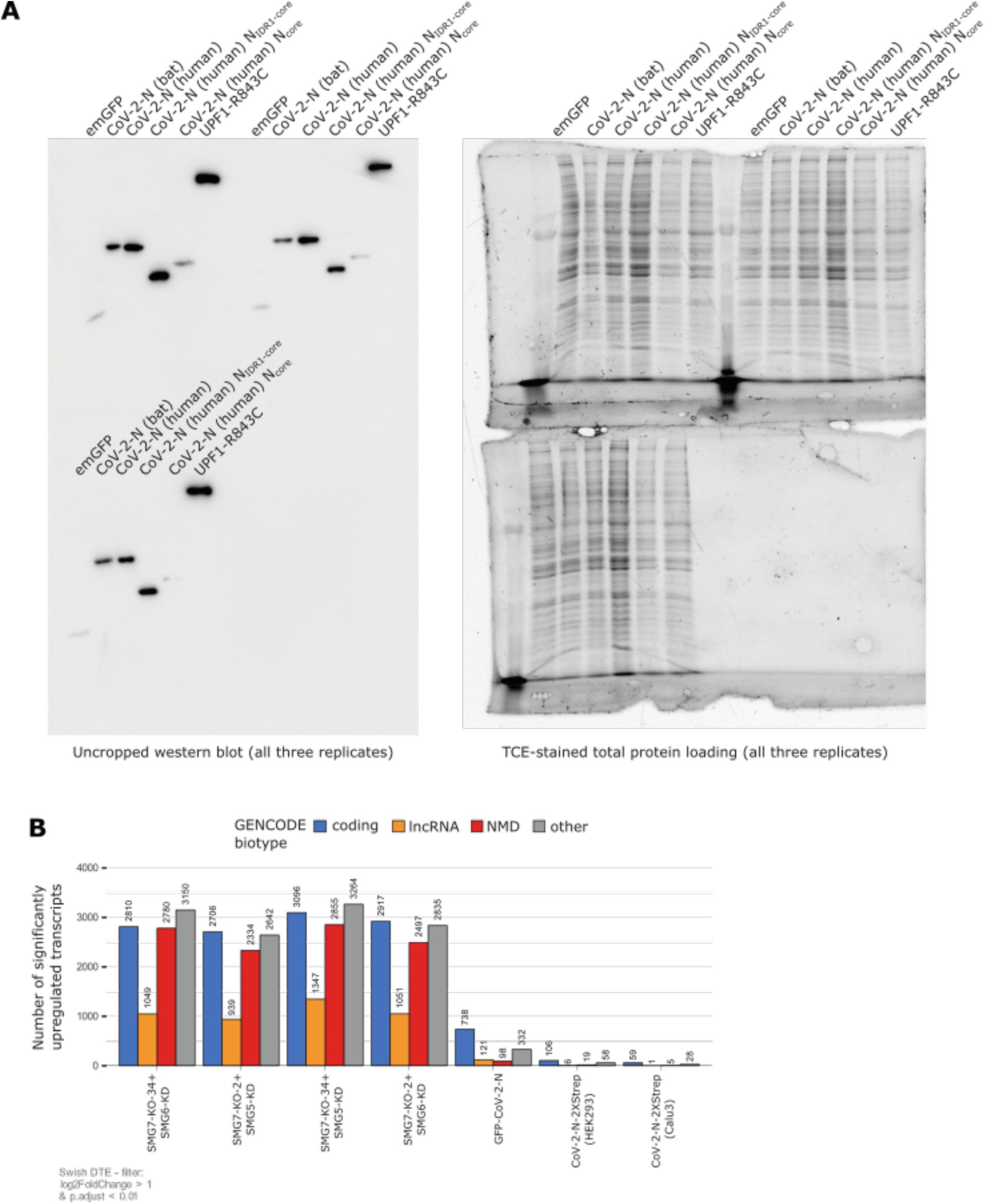
**A)** Uncropped western blots corresponding to Figure 5B (left) and total protein loading of SDS-PAA gels using TCE staining (right). **B)** Absolute numbers of significantly upregulated transcripts per GENCODE biotype (cut-offs are indicated).

**Supplementary Table 1.**
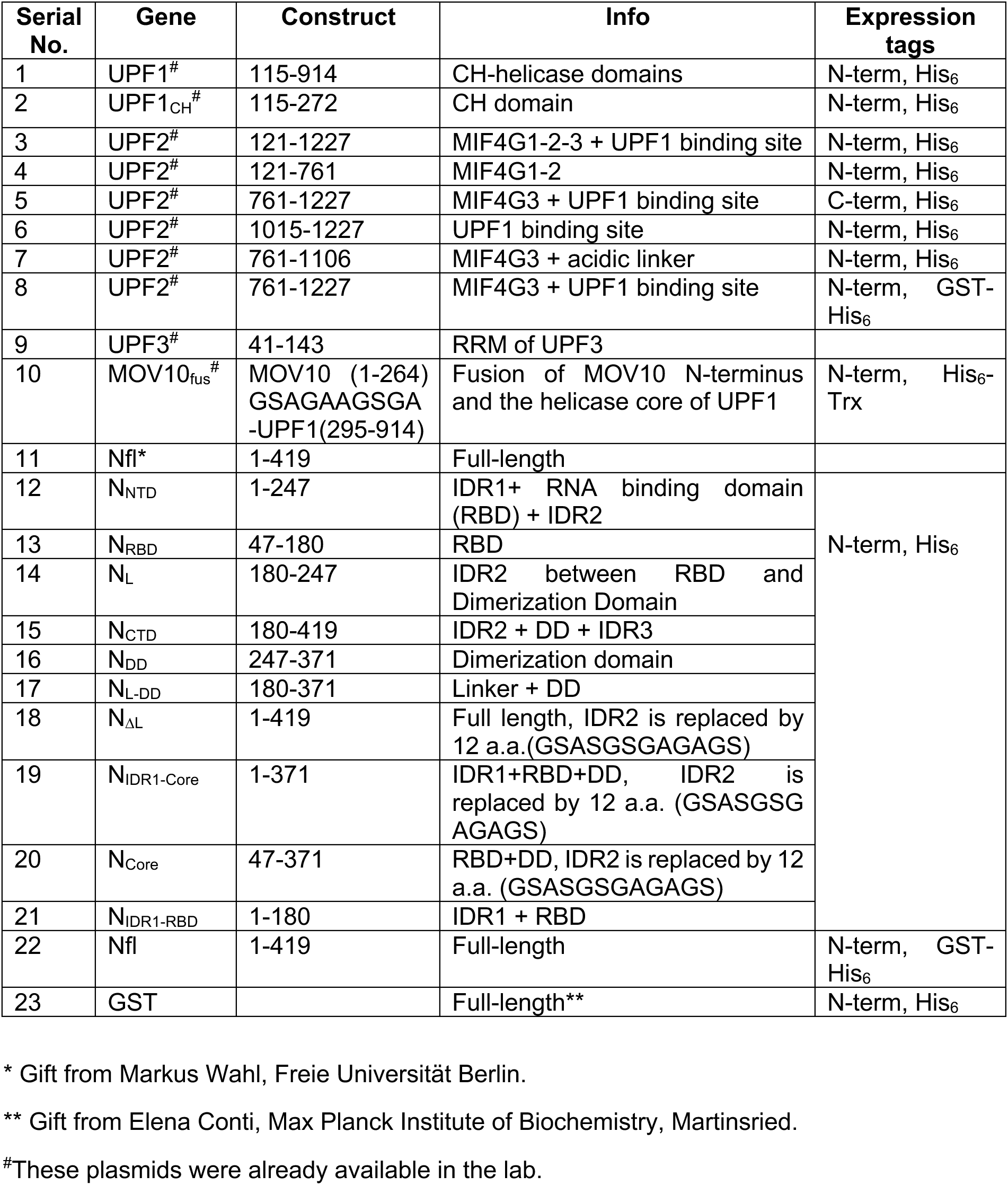
Plasmids used in this study. *E. coli* expression (Vector backbone for all plasmids: pET28)

**Supplementary Table 2.**
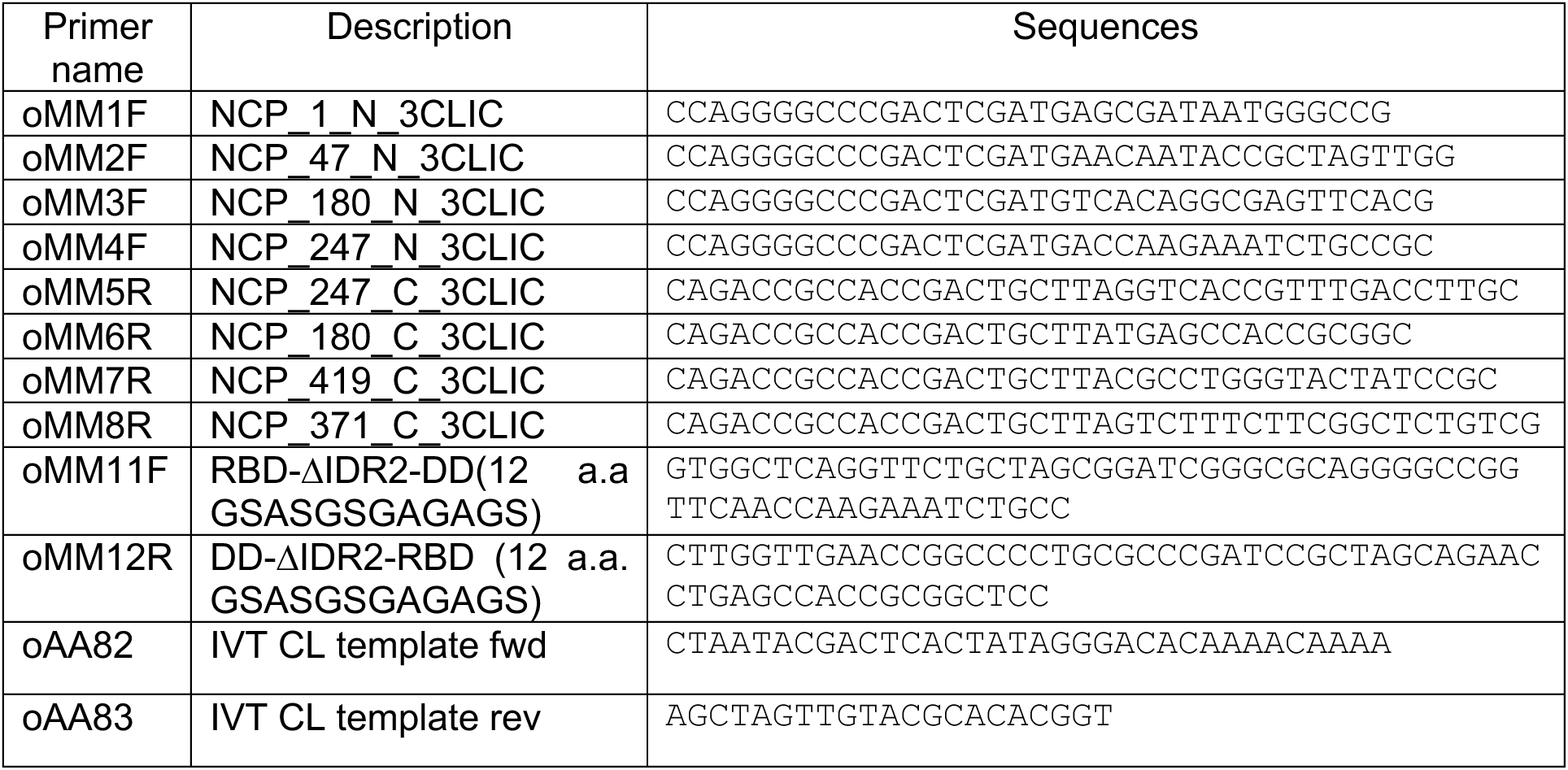
Primers used in this study.

