## Supplementary figure for "Functional implications of the interaction of the SARS-CoV-2 Nucleocapsid protein with factors involved in nonsense-mediated mRNA decay"

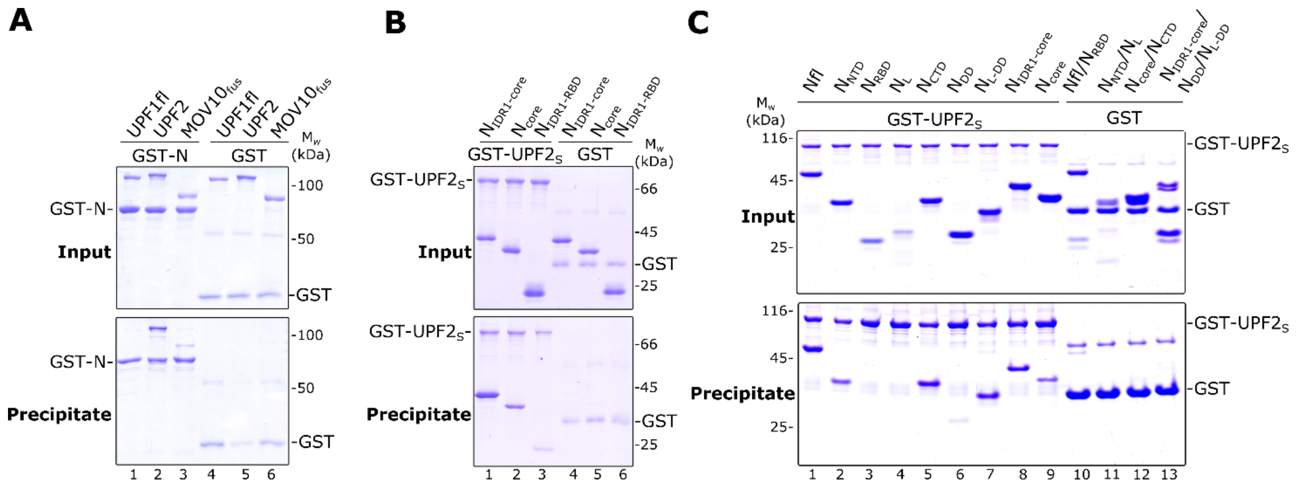


**D**

```
sp|P03416|NCAP_CVMA5      MSFVPGQENAGGRSSSVNRAGNGILKKTTWADQTERGPNNQNR----GRRNQPKQTATTQ     56
sp|K9N4V7|NCAP_MERS1      ------------M------ASPAAPRAVSFADNNDITNTN----LSRGRGRNPK------     32
sp|P59595|NCAP_SARS       MS------DNGPQ------SNQRSAPRITFGGPTDSTDNNQNGGRNGARPKQRR------     42
sp|P0DTC9|NCAP_SARS2      MS------DNGPQ-------NQRNAPRITFGGPSDSTGSNQNGERSGARSKQRR------     41
                                              .       ::.. .:   .*       .* .: :

sp|P03416|NCAP_CVMA5      PNSGSVVPHYSWFSGITQFQKGKEFQFAEGQGVPIANGIPASEQKGYWYRHNRRSFKTPD     116
sp|K9N4V7|NCAP_MERS1      -PRAAPNNTVSWYTGLTQHGK-VPLTFPPGQGVPLNANSTPAQNAGYWRRQDR-KINTGN     89
sp|P59595|NCAP_SARS       -PQGLPNNTASWFTALTQHGK-EELRFPRGQGVPINTNSGPDDQIGYYRRATR-RVRGGD     99
sp|P0DTC9|NCAP_SARS2      -PQGLPNNTASWFTALTQHGK-EDLKFPRGQGVPINTNSSPDDQIGYYRRATR-RIRGGD     98
                             .      **::.:**. *   : *  *****:  .    ::  **: *  *  ..  :
                                             RBD

sp|P03416|NCAP_CVMA5      GQQKQLLPRWYFYYLGTGPHAGASYGDSIEGVFWVANSQADTNTRSDIVERDPSSHEAIP     176
sp|K9N4V7|NCAP_MERS1      G-IKQLAPRWYFYYTGTGPEAALPFRAVKDGIVWVHEDGATDAPS-TFGTRNPNNDSAIV     147
sp|P59595|NCAP_SARS       GKMKELSPRWYFYYLGTGPEASLPYGANKEGIVWVATEGALNTPKDHIGTRNPNNNAATV     159
sp|P0DTC9|NCAP_SARS2      GKMKDLSPRWYFYYLGTGPEAGLPYGANKDGIIWVATEGALNTPKDHIGTRNPANNAAIV     158
                          *  *:* ******* ****.*.  :    :*:.**  . *       :  *:* .. *
                                             RBD

sp|P03416|NCAP_CVMA5      TRFAPGTVLPQGFYVEGSGRSAPAS-RSGSRSQSRGPNNRARSSSNQRQPASTV-KPD-M     233
sp|K9N4V7|NCAP_MERS1      TQFAPGTKLPKNFHIEGTGGNSQSSSRASSVSRNSSRSSSQGS-RSGNSTRGTSPGPSGI     206
sp|P59595|NCAP_SARS       LQLPQGTTLPKGFYAEGSRGGSQASSRSSSRSRGNSRNSTPGS-SRGNSPARMA---SGG     215
sp|P0DTC9|NCAP_SARS2      LQLPQGTTLPKGFYAEGSRGGSQASSRSSSRSRNSSRNSTPGS-SRGTSPARMA---GNG     214
                           ::  ** **:.*: **:  .: :* *:.* *:. . ..   *     .        .
                               RBD

sp|P03416|NCAP_CVMA5      -AEEIAALVLAKLG--------K-DAGQPKQVTKQSAKEVRQKILNKPRQKRTPNKQCPV     283
sp|K9N4V7|NCAP_MERS1      GAVGGDLLYLDLLNRLQALESGKVKQSQPKVITKKDA----AAAKNKMRHKRTSTKSFNM     262
sp|P59595|NCAP_SARS       GETALALLLLDRLNQLESKVSGKGQQQQGQTVTKKSA----AEASKKPRQKRTATKQYNV     271
sp|P0DTC9|NCAP_SARS2      GDAALALLLLDRLNQLESKMSGKGQQQQGQTVTKKSA----AEASKKPRQKRTATKAYNV     270
                                 * *  *.        * .  * : :**:.*        :* *:*** .*   :
                                                                   DD

sp|P03416|NCAP_CVMA5      QQCFGKRGPNQ---NFGGSEMLKLGTSDPQFPILAELAPTVGAFFFGSKLELVKKNSGGA     340
sp|K9N4V7|NCAP_MERS1      VQAFGLRGPGDLQGNFGDLQLNKLGTEDPRWPQIAELAPTASAFMGMSQFKLTHQNN---     319
sp|P59595|NCAP_SARS       TQAFGRRGPEQTQGNFGDQDLIRQGTDYKHWPQIAQFAPSASAFFGMSRIGMEVTP----     327
sp|P0DTC9|NCAP_SARS2      TQAFGRRGPEQTQGNFGDQELIRQGTDYKHWPQIAQFAPSASAFFGMSRIGMEVTP----     326
                           *.** *** :   ***. :: : **.  ::* :*::**:..**:  *:: :
                                                     DD

sp|P03416|NCAP_CVMA5      DEPTKDVYELQYSGAVRFDSTLPGFETIMKVLNENLNAYQKDGGADVVSPKPQRKGRRQA     400
sp|K9N4V7|NCAP_MERS1      DDHGNPVYFLRYSGAIKLDPKNPNYNKWLELLEQNIDAYKTFPKKEKKQKAPKEESTDQM     379
sp|P59595|NCAP_SARS       -----SGTWLTYHGAIKLDDKDPQFKDNVILLNKHIDAYKTFPPTEPKKDKKKKTDEA--     380
sp|P0DTC9|NCAP_SARS2      -----SGTWLTYTGAIKLDDKDPNFKDQVILLNKHIDAYKTFPPTEPKKDKKKKADET--     379
                                   * * **:::* . * ::  : :*:::::**:.    :  .   :. .
                                                  DD

sp|P03416|NCAP_CVMA5      QEKK--DEVDNVSVAKPKSSVQRNVSRELTPEDRSLLAQILDDGVVPDGLEDDSNV         454
sp|K9N4V7|NCAP_MERS1      SEPPKEHRVQGTQRTRTRPSVQPGPMIDVNTD------------------------         411
sp|P59595|NCAP_SARS       QPLPQ--------RQKKQPTVTLLPAADMDDFSRQLQNSMS--GASADSTQ----A         422
sp|P0DTC9|NCAP_SARS2      QALPQ--------RQKKQQTVTLLPAADLDDFSKQLQQSMS--SADSTQA------         419
                          .              : : :*      ::
```

**Figure S1. The SARS-CoV-2 Nucleocapsid (N) protein directly interacts with the core NMD factor UPF2. A)** GST-pulldown assay of full length UPF1-(UPF1fl), UPF2, and $MOV10_{fus}$ (a fusion of the N-terminus of MOV10 with the helicase core of UPF1) with full-length GST-N as a bait. GST was used as a negative control in all such assays. The top and bottom panel depict inputs and precipitates, respectively, in this and all the other GST-pull down assays. GST-N binds to UPF2 but not UPF1fl or $MOV10_{fus}$. **B)** GST-pulldown assay to compare binding of the IDRs and the structured domains of N to UPF2. Here GST-$UPF2_S$ is used as a bait and $N_{IDR1\text{-}core}$, $N_{core}$ and $N_{IDR1\text{-}RBD}$ as preys. Variants of N containing only one of the three identified binding motifs (IDR1, linker and DD), such as $N_{core}$ and $N_{IDR1\text{-}RBD}$, show weaker binding to UPF2 in comparison to $N_{IDR1\text{-}core}$, which contains two of three binding motifs. We conclude that no single binding site of N can mediate a strong interaction with UPF2: a combination of any two binding sites are necessary for a stable N-UPF2 interaction. **C)** A complete gel corresponding to Figure 1D, including negative controls using GST as a bait. Domain organization of the N-constructs shown in Figure 1A. **D)** Multiple sequence alignment using Clustal Omega (Clustal O 1.2.4) with N-protein sequences from different coronaviruses (P03416: Murine Hepatitis Virus, K9N4V7: MERS, P59595: SARS-CoV and P0DTC9: SARS-CoV-2 strain, acquired from Uniprot: https://www.uniprot.org/). The sequence similarities across species are indicated as follows: * - fully conserved, : - strongly similar, . – weakly similar. The domain organization of SARS-CoV2 N is indicated below the alignment. Shaded boxes denote structured domains and dashed lines depict intrinsically disordered regions (IDRs) flanking the domains. Sequence conservation is highest among amino acids within the structured regions and considerably lower across the IDRs.

**A**

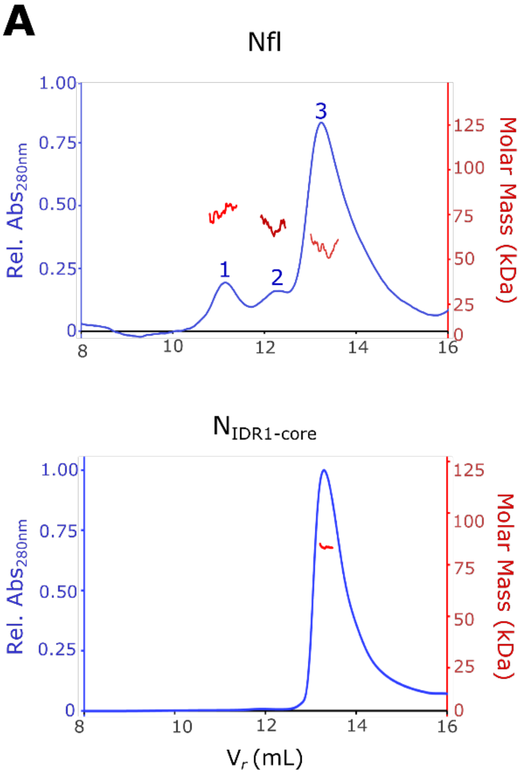


| Theoretical $M_w$ (kDa) | Experimental $M_w$ (kDa) |
|---|---|
| Nfl<br>Monomer = 45.6<br>Dimer = 91.2 | Peak 1 = 80.1 ± 9.2<br>Peak 2 = 72.8 ± 9.2<br>Peak 3 = 58.3± 5.7 |
| $N_{IDR1\text{-}core}$<br>Monomer = 36.7<br>Dimer = 73.4 | Peak = 83.6 ± 1.5 |

**B**

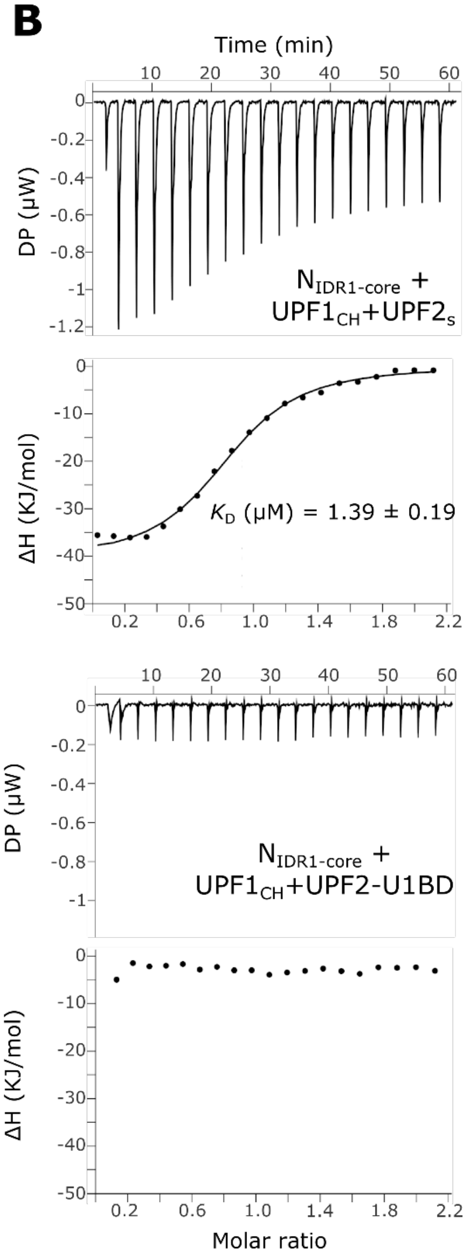


**Figure S2. A)** Size Exclusion Chromatography-Multi-angle Light Scattering (SEC-MALS) analysis to determine the oligomerization state of full-length N (Nfl) and $N_{IDR1\text{-}core}$ and the molar mass (kDa) across the observed peaks. The experimentally-obtained values are indicated in the table below, along with the theoretical molar mass (calculated using the Expasy Protparam tool) for distinct oligomerization states of each N construct. Nfl exists as a mixture of indistinct oligomers in solution whereas $N_{IDR1\text{-}core}$, lacking the long inter-domain linker, forms a homogenous dimer. **B)** Isothermal titration calorimetry (ITC) experiments of $N_{IDR1\text{-}core}$ with complexes of $UPF1_{CH}$ and UPF2 ($UPF2_S$ and UPF2-U1BD, top and bottom panels, respectively) show that addition of UPF1 does not influence the binding affinity of N to UPF2 (compare $K_D$ derived from this experiment to that in Figure 2A).

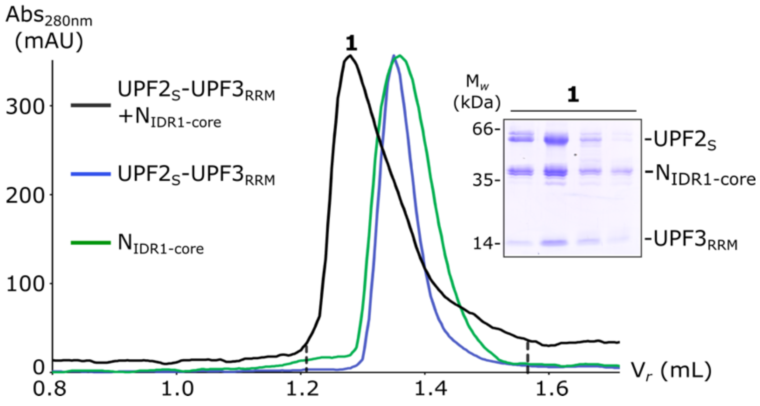


**Figure S3.** Analytical size-exclusion chromatography (SEC) shows that UPF2 binds N and UPF3 simultaneously. SDS-PAGE analysis corresponding to peak 1 of the black trace depicts formation of a ternary complex of $UPF2_S$-$UPF3_{RRM}$-$N_{IDR1\text{-}core}$. The terms $Abs_{280nm}$ and $V_r$ refer to absorbance at 280 nm and retention volume, respectively.

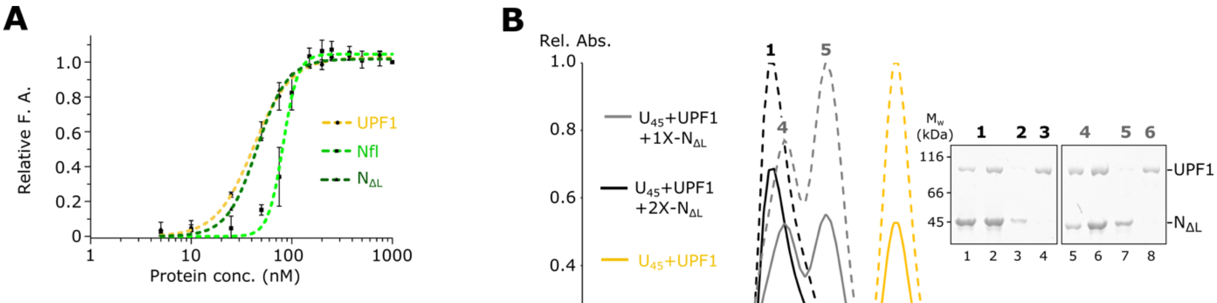


| Protein | $K_D$ (nM) |
|---|---|
| UPF1 | 41.9 ± 0.9 |
| Nfl | 82.1 ± 1.8 |
| $N_{\Delta L}$ | 45.5 ± 1.1 |

**Figure S4. A)** Fluorescence anisotropy measurements using a 5'-end 6-FAM labeled12-mer poly-U RNA ($U_{12}$) shows that UPF1 (in yellow trace), Nfl (light green) and $N_{\Delta L}$ (dark green) have comparable binding affinities towards the RNA with a $K_D$ of ~ 50 nM for UPF1-RNA, 82 nM for Nfl-RNA and 45 nm for $N_{\Delta L}$-RNA. The data points and their error bars represent the mean of fluorescence anisotropy (calculated using Graph Pad Prism 5.00) and the standard deviation of independent experiments. **B)** Analytical size exclusion chromatography (SEC) and the corresponding SDS-PAGE analyses of mixtures of UPF1, a 45-mer poly-U RNA ($U_{45}$) and $N_{\Delta L}$, where UPF1 and $U_{45}$ are in equi-molar amounts and $N_{\Delta L}$ is added either in equi-molar amounts to UPF1 (grey trace) and in 2-fold excess to UPF1 (black trace). Although N and UPF1 can co-occupy the RNA (lanes 1 and 5, corresponding to peaks 1 and 4), a small peak corresponding to RNA-free UPF1 is always observed (lanes 4 and 8, corresponding to peaks 3 and 6). Addition of excess N leads to an increase in free UPF1 (compare peaks 3 and 6). Solid and dashed lines refer to absorbance at 280 nm and 260 nm, respectively.

**A**

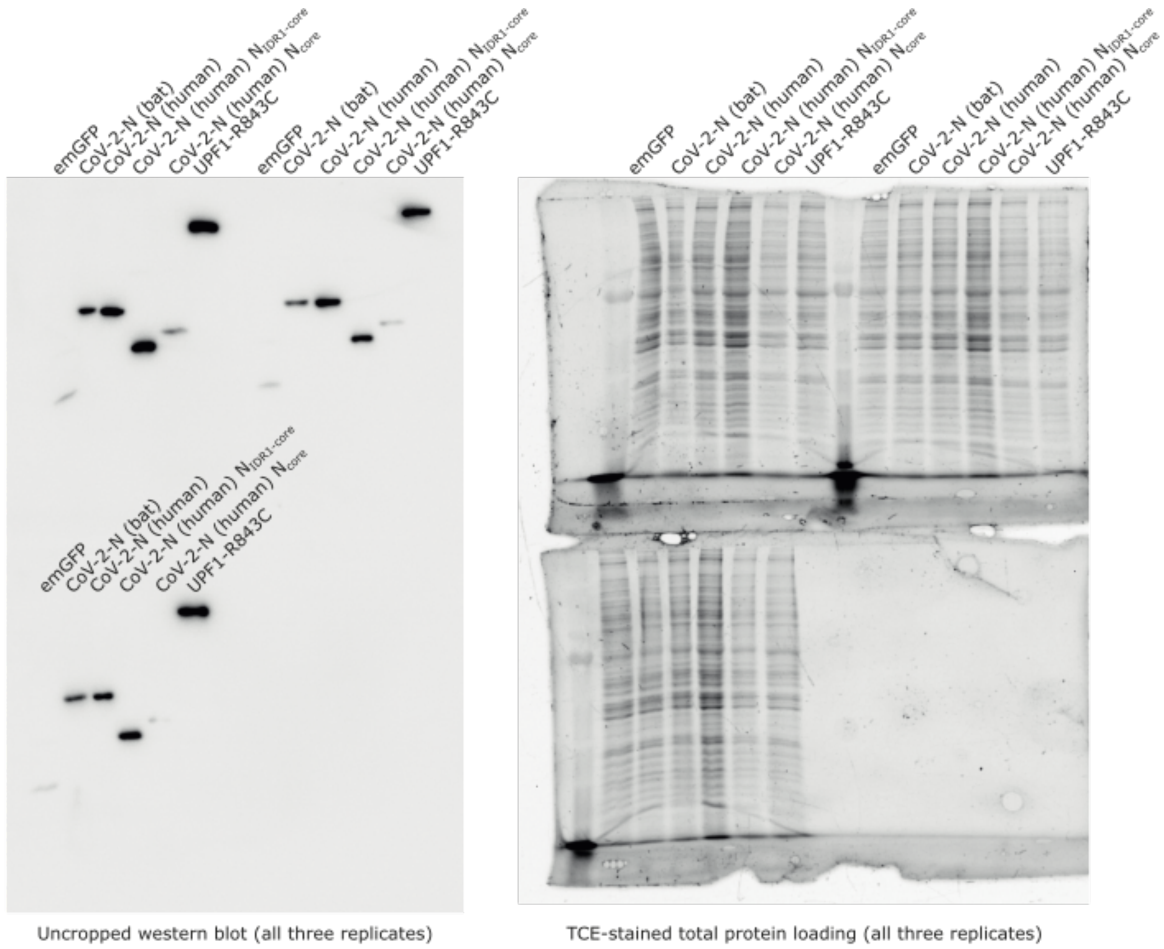


**B**

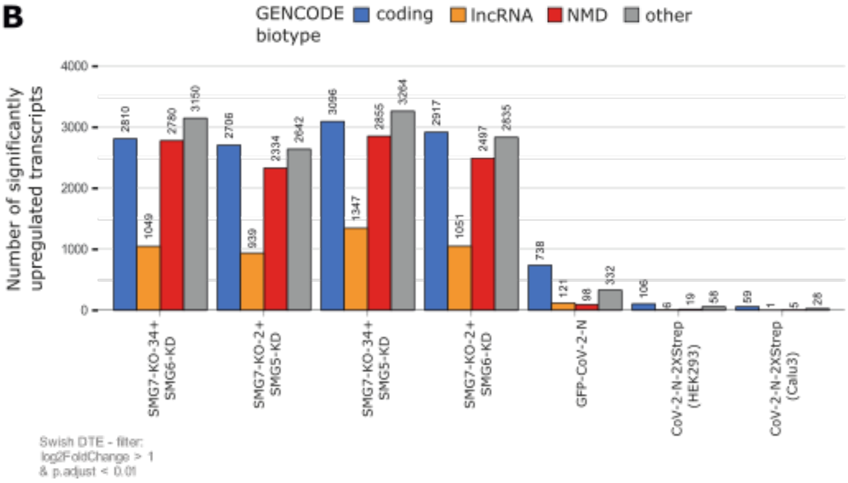


**Figure S5 A)** Uncropped western blots corresponding to Figure 5B (left) and total protein loading of SDS-PAA gels using TCE staining (right). **B)** Absolute numbers of significantly upregulated transcripts per GENCODE biotype (cut-offs are indicated).

| **Composition of master mix** | **Volume** | **Final Conc.** |
|---|---|---|
| 5x Unwinding buffer | 8 µl | 1x |
| 100 mM magnesium acetate | 0.8 µl | 2 mM |
| 100 mM DTT (freshly prepared) | 0.8 µl | 2 mM |
| Duplex (RNA-DNA) | 5.5 µl | 75 nM |
| Nuclease-free water | 1.4 µl | |

**Supplementary Table 1. Plasmids used in this study**

*E. coli* expression (Vector backbone for all plasmids: pET28)

| Serial No. | Gene | Construct | Info | Expression tags |
|---|---|---|---|---|
| 1 | UPF1[#] | 115-914 | CH-helicase domains | N-term, His$_6$ |
| 2 | UPF1$_{CH}$[#] | 115-272 | CH domain | N-term, His$_6$ |
| 3 | UPF2[#] | 121-1227 | MIF4G1-2-3 + UPF1 binding site | N-term, His$_6$ |
| 4 | UPF2[#] | 121-761 | MIF4G1-2 | N-term, His$_6$ |
| 5 | UPF2[#] | 761-1227 | MIF4G3 + UPF1 binding site | C-term, His$_6$ |
| 6 | UPF2[#] | 1015-1227 | UPF1 binding site | N-term, His$_6$ |
| 7 | UPF2[#] | 761-1106 | MIF4G3 + acidic linker | N-term, His$_6$ |
| 8 | UPF2[#] | 761-1227 | MIF4G3 + UPF1 binding site | N-term, GST-His$_6$ |
| 9 | UPF3[#] | 41-143 | RRM of UPF3 | |
| 10 | MOV10$_{fus}$[#] | MOV10 (1-264) GSAGAAGSGA -UPF1(295-914) | Fusion of MOV10 N-terminus and the helicase core of UPF1 | N-term, His$_6$-Trx |
| 11 | Nfl* | 1-419 | Full-length | |
| 12 | N$_{NTD}$ | 1-247 | IDR1+ RNA binding domain (RBD) + IDR2 | |
| 13 | N$_{RBD}$ | 47-180 | RBD | N-term, His$_6$ |
| 14 | N$_{L}$ | 180-247 | IDR2 between RBD and Dimerization Domain | |
| 15 | N$_{CTD}$ | 180-419 | IDR2 + DD + IDR3 | |
| 16 | N$_{DD}$ | 247-371 | Dimerization domain | |
| 17 | N$_{L-DD}$ | 180-371 | Linker + DD | |
| 18 | N$_{\Delta L}$ | 1-419 | Full length, IDR2 is replaced by 12 a.a.(GSASGSGAGAGS) | |
| 19 | N$_{IDR1-Core}$ | 1-371 | IDR1+RBD+DD, IDR2 is replaced by 12 a.a. (GSASGSG AGAGS) | |
| 20 | N$_{Core}$ | 47-371 | RBD+DD, IDR2 is replaced by 12 a.a. (GSASGSGAGAGS) | |
| 21 | N$_{IDR1-RBD}$ | 1-180 | IDR1 + RBD | |
| 22 | Nfl | 1-419 | Full-length | N-term, GST-His$_6$ |
| 23 | GST | | Full-length** | N-term, His$_6$ |

* Gift from Markus Wahl, Freie Universität Berlin.

** Gift from Elena Conti, Max Planck Institute of Biochemistry, Martinsried.

[#]These plasmids were already available in the lab.

**Supplementary Table 2. Primers used in this study**

| Primer name | Description | Sequences |
|---|---|---|
| oMM1F | NCP_1_N_3CLIC | CCAGGGGCCCGACTCGATGAGCGATAATGGGCCG |
| oMM2F | NCP_47_N_3CLIC | CCAGGGGCCCGACTCGATGAACAATACCGCTAGTTGG |
| oMM3F | NCP_180_N_3CLIC | CCAGGGGCCCGACTCGATGTCACAGGCGAGTTCACG |
| oMM4F | NCP_247_N_3CLIC | CCAGGGGCCCGACTCGATGACCAAGAAATCTGCCGC |
| oMM5R | NCP_247_C_3CLIC | CAGACCGCCACCGACTGCTTAGGTCACCGTTTGACCTTGC |
| oMM6R | NCP_180_C_3CLIC | CAGACCGCCACCGACTGCTTATGAGCCACCGCGGC |
| oMM7R | NCP_419_C_3CLIC | CAGACCGCCACCGACTGCTTACGCCTGGGTACTATCCGC |
| oMM8R | NCP_371_C_3CLIC | CAGACCGCCACCGACTGCTTAGTCTTTCTTCGGCTCTGTCG |
| oMM11F | RBD-ΔIDR2-DD(12 a.a GSASGSGAGAGS) | GTGGCTCAGGTTCTGCTAGCGGATCGGGCGCAGGGGCCGG TTCAACCAAGAAATCTGCC |
| oMM12R | DD-ΔIDR2-RBD (12 a.a. GSASGSGAGAGS) | CTTGGTTGAACCGGCCCCTGCGCCCGATCCGCTAGCAGAAC CTGAGCCACCGCGGCTCC |
| oAA82 | IVT CL template fwd | CTAATACGACTCACTATAGGGACACAAAACAAAA |
| oAA83 | IVT CL template rev | AGCTAGTTGTACGCACACGGT |
